# Extrachromosomal Circular DNA Function as Mobile Elements to Alter the Mammalian Germline Genome

**DOI:** 10.64898/2026.09.03.749254

**Authors:** Xin Zhang, Melanie Evans, Shreya Rajachandran, Angie Huang, Yanfeng Zhang, Karla Saner, Lin Xu, Kyle E. Orwig, Remus-Laurentiu Stana, Orhan Bukulmez, Vladimir Seplyarskiy, Haiqi Chen

## Abstract

The viability of any species including the human requires that the germline genome is kept stable as it is transmitted across generations by the germ cells. Failure to safeguard the genome integrity and stability would lead to inherited diseases and infertility. Thus, a better understanding of the mechanisms that alter the germline genome is crucial to ensure human health and our continuation as a species. Here, we show that the extrachromosomal circular DNA (eccDNA) in the mouse and human male germline represents a new mechanism in altering the mammalian germline genome. To enable the tracking of germline eccDNA *in vivo*, we established a novel mouse model that allows the generation of a reporter eccDNA in a cell type-specific manner. Using this mouse model, we showed that eccDNA formed in the developing male germ cells can integrate into the germline genome. Using eccDNA-containing sperm for *in vitro* fertilization led to the eccDNA sequence being inherited by the embryos. By analyzing a large cohort of long-read whole genome sequencing data, we showed that eccDNA-mediated germline genome insertions represent an important source of human genome structural variations. Finally, by leveraging human sperm samples, we found that diabetes induces an increase in sperm eccDNA quantity, which is mediated at least in part through poly (ADP-ribose) polymerases. Together, our results provide new insights into how the mammalian germline genome can be altered, with important implications for human health and genome evolution.

## Introduction

The integrity of the mammalian germline genome is essential to produce viable gametes and for successful reproduction [1, 2]. Yet, mechanisms exist to alter the germline genome, such as transposable elements (TEs)-mediated genome insertions [3, 4] and aging-induced de novo mutations [5, 6]. These changes to the germline genome may not only impact the organism itself, in terms of reproductive success, but also for subsequent generations, in terms of their development. Thus, a better of understanding of the mechanisms that alter the germline genome is crucial to ensure reproductive health and our continuation as a species.

Extrachromosomal circular DNA (eccDNA) originate from the self-circularization of linear genome fragments and can be found in various species and cell types including the mouse and human sperm [7–10]. However, the functional role and the biogenesis mechanism of germline eccDNA remain largely unexplored. Recent studies on somatic cells show that circular DNA may insert into the linear genome. For example, one study of domestic cattle showed that a ∼500 kilo base pair (kbp) genome fragment deleted from chromosome 29 can circularize and then re-integrate into a region on chromosome 6 to affect pigmentation patterns [11]. In cancer cells, circular DNA can re-integrate into the linear cancer genome, causing intra- and inter-chromosomal rearrangements [12, 13]. Thus, we hypothesized that eccDNA can alter the germline genome landscape.

Studying the mammalian germline eccDNA faces the following challenges. First, unlike cancer circular DNA which are often 100 kb to several Mb in size and are highly amplified (tens to hundreds of copies per cell) [14, 15], germline eccDNA are 50 kb or less and are often at a single copy per cell [7], making it difficult to identify, target, and track germline eccDNA. Second, almost the entire germline genome can give rise to eccDNA [7, 16], making the sequence identities of the germline eccDNA in an individual unpredictable. Third, the reproducible rate of germline eccDNA between biological replicates is low [16], meaning that different individuals may have completely different repertoire of germline eccDNA. Lastly, a lack of in vitro models that faithfully recapitulate the in vivo developmental process of the mammalian germ cells especially for the human makes it even more challenging to track the behavior of germline eccDNA during germ cell development and across generations. Therefore, significant technical innovations are needed to study the function and biogenesis of mammalian germline eccDNA.

Here, we established a reporter eccDNA mouse model to allow for the tracking of germline eccDNA in vivo. Leveraging short- and long-read sequencing and various molecular and cellular assays, we showed that male germline eccDNA can induce alterations to the germline genome both in the mouse and the human. We also showed that poly (ADP-ribose) polymerases may regulate the biogenesis of germline eccDNA.

## Results

**A genetic mouse model to track male germline eccDNA.**

To test our hypothesis that eccDNA can alter the mammalian germline genome, we first sought to track the fate of germline eccDNA in vivo. Due to the ethical and practical constraints of human germline research, we set out to engineer a mouse model of male germline eccDNA given that mouse male germ cells also harbor eccDNA endogenously [10, 16]. We designed the mouse model to produce reporter eccDNA in the male germline using the Cre-loxP system. Specifically, an eccDNA cassette was generated by flanking a chicken β-actin (CAG) promoter-controlled EGFP sequence in the inverted configuration with two loxP sites (**Figure 1A**). This cassette, when at its linear form, cannot drive EGFP expression. Upon Cre recombinase-mediated excision, however, the eccDNA cassette self-circularizes to form a reporter eccDNA that would drive the expression of EGFP (**Figure 1A**). The EGFP expression would allow cells that contain the reporter eccDNA to be identified and tracked. We hereafter refer to this reporter eccDNA as eccDNA^EGFP^.

**Figure 1.**
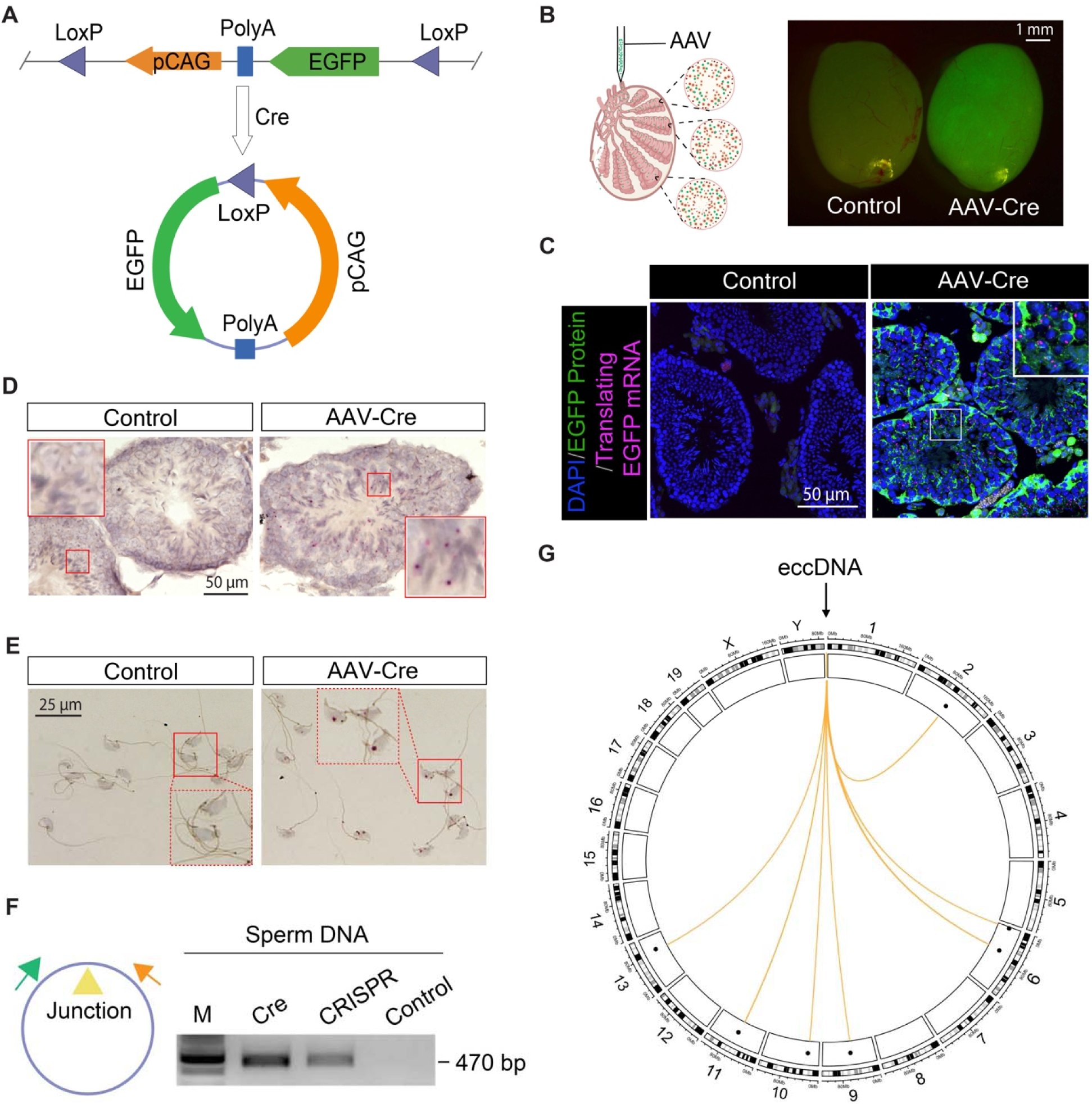
Germline eccDNA can re-integrate into the linear genome. (A) Design of the eccDNA cassette. (B) Rete testis injections of the eccDNA mouse with AAV-Cre led to EGFP expression in the seminiferous tubules. (C) Ribosome-associated EGFP mRNA and EGFP protein were observed in the seminiferous epithelium of eccDNA mice injected with AAV-Cre. (D) DNA ISH showed positive signals (red loci) of eccDNA^EGFP^ in the seminiferous epithelium of eccDNA mice injected with AAV-Cre. (E) DNA ISH showed positive signals (red loci) of eccDNA^EGFP^ in the cauda epidydimal sperm of eccDNA mice injected with AAV-Cre. (F) Positive signals of the eccDNA^EGFP^ junction region were observed in PCR using sperm DNA from eccDNA mice either injected with AAV-Cre or targeting CRISPR-C. M, marker. (G) Visualization of eccDNA^EGFP^ insertion events in cauda epidydimal sperm of eccDNA mice injected with AAV-Cre.

We first tested this design by transfecting the HEK293T cells with plasmids containing the eccDNA cassette and the Cre recombinase, respectively. Encouragingly, the EGFP expression was only observed in the presence of both the eccDNA cassette and the Cre recombinase (**Figure S1A**), suggesting the successful generation of eccDNA^EGFP^. To enable the generation of the eccDNA^EGFP^ in vivo, we established a mouse line by stably integrating the eccDNA cassette into the ROSA26 safe harbor locus of the mouse genome (hereafter referred as the eccDNA mouse). To induce the eccDNA^EGFP^ formation in the developing male germ cells, we injected the Cre recombinase-expressing adeno-associated viruses (AAV-Cre) into the testis of a 3-week-old eccDNA mouse through the rete testis (**Figure 1B**). This rete testis injection method introduces AAV particles into the seminiferous tubules, enabling the transduction of developing male germ cells at various developmental stages (**Figure S1B**). 4 weeks after AAV-Cre injection, the whole-mount injected testes of eccDNA mice were imaged using a stereoscope and green fluorescence signals were observed in the AAV-Cre-treated group but not in the Cre-negative control group (**Figure 1B**), consistent with the in vitro cell culture data. Imaging of the cross-sections of the injected testes showed green fluorescence signals in both Sertoli cells and the developing germ cells in the AAV-Cre-treated group but not in the Cre-negative control group (**Figure 1C**). Using a RIBOmap-based RNA in situ hybridization approach (Methods), we detected ribosome-associated EGFP mRNA in the AAV-Cre-injected testis cross-sections (**Figure 1C**), suggesting that the green fluorescence signals of the AAV-Cre-injected testis came from EGFP. To further confirm the successful generation of eccDNA^EGFP^ *in vivo*, we performed chromogenic DNA in situ hybridization (DNA ISH) using probes that specifically target the junction region of eccDNA^EGFP^ (Methods). The positive signals (red foci) were only found in the seminiferous epithelia in the AAV-Cre-treated group but not in the Cre-negative control group (**Figure 1D**).

To rule out the possibility that the EGFP signals might be caused by abnormal Cre activity-induced genome rearrangements, we used an alternative approach called CRISPR-C [17] to induce eccDNA^EGFP^ formation in vivo. In this experiment, we injected into the testes of the eccDNA mice with AAVs that express both SaCas9 protein (a compact variant of the Cas9 family protein) and a pair of gRNAs. The gRNA pair specifically targets the two ends of the integrated eccDNA cassette at the ROSA26 locus, leading to the deletion and subsequent circularization of the eccDNA cassette (**Figure S1C**). A pair of non-targeting gRNAs was used as a negative control. As expected, the green fluorescence signals were observed in the testis from the targeting gRNA group but not from the non-targeting gRNA group (**Figure S1D**).

Together, these data demonstrate that we have successfully generated an eccDNA reporter mouse model that enables the production of eccDNA *in vivo*.

## Male germline eccDNA can integrate into the linear mouse genome

After establishing the eccDNA mouse model, we went on to track the fate of eccDNA^EGFP^ generated in developing male germ cells. To this end, we first collected the cauda epidydimal sperm from the eccDNA mice 4 weeks after the injection of AAVs and performed DNA ISH using probes that target the junction region of eccDNA^EGFP^. We observed positive DNA ISH signals only in the AAV-Cre-treated group but not in the Cre-negative control group (**Figure 1E**), suggesting the presence of eccDNA^EGFP^ in the sperm of AAV-Cre-injected eccDNA mice. Next, we collected the DNA of these sperm and treated the DNA with exonuclease to digest the linear DNA. The remaining circular DNA was used for PCR with primers that correspond to the flanking regions of the eccDNA^EGFP^ junction. After performing electrophoresis using the PCR products, we saw a specific band corresponding to the correct size of the eccDNA^EGFP^ junction region whose sequence identity was confirmed by Sanger sequencing (**Figure 1F**). Using the same approach, we also confirmed the sequence identity of the eccDNA^EGFP^ extracted from eccDNA mice that were subject to the targeting CRISPR-C (**Figure 1F**). Together, these data suggest that at least a portion of eccDNA generated in developing male germ cells can remain as extrachromosomal elements in sperm.

Next, to test if eccDNA^EGFP^ can re-integrate into the linear genome, we collected approximately one million cauda epidydimal sperm from an eccDNA mouse 4 weeks after the AAV-Cre injection, extracted sperm total DNA, and performed Nanopore long-read whole genome sequencing (no PCR amplification was performed). Encouragingly, despite sequencing the bulk sample at only 44x single-copy genome coverage, we observed one instance where the eccDNA^EGFP^ sequence was found inserted into a site at the chromosome 1 (**Figure S2A**). This insertion site is different from the original integration site of the eccDNA reporter cassette which is at the ROSA26 locus of the chromosome 6, suggesting the re-integration of the eccDNA^EGFP^ into a different part of the linear germline genome.

To identify additional eccDNA^EGFP^-mediated germline genome insertion events, we developed a more sensitive, targeted approach (Methods). This targeted approach enriches for chimeric sequences that contain both the eccDNA^EGFP^ sequence and the genomic sequence using Tn5 transposase-mediated random genome tagmentation followed by nested PCR (**Figure S2B**). The PCR products are then subject to Nanopore long-read sequencing to identify potential eccDNA^EGFP^ genome insertion events. To avoid PCR artifacts, only insertion sites that are supported by at least two independent, unique sequencing reads are retained. We applied this targeted long-read sequencing approach to a bulk population of approximately one million sperm from an AAV-Cre-injected eccDNA mouse and identified 7 eccDNA^EGFP^-mediated germline genome insertion events (**Figure 1G**). Although 6 out of 7 insertions took place in distal intergenic regions and introns of the mouse genome, one insertion was found at the promoter of a gene called *Gm10800* (also known as *mPIRO-1* whose function may be related to bone homeostasis [18]) (**Figure S2C**), suggesting a potential disruption of the germline genome function.

To rule out the possibility that these genome insertion events were caused by abnormal activities of the Cre recombinase, we performed the same targeted Nanopore sequencing experiment on sperm DNA from eccDNA mouse testes treated with targeting CRISPR-C. Reassuringly, we also observed an instance of eccDNA^EGFP^ insertion in the germline genome (**Figure S2D**). The reason we did not detect more eccDNA^EGFP^ insertion events could be due to a lower genome cutting efficiency of the CRISPR-C approach to generate eccDNA^EGFP^ compared to the Cre-mediated approach.

Together, our data suggest that eccDNA generated in the developing male germ cells can either persist as extrachromosomal genetic elements or re-integrate into the germline genome.

## Germline eccDNA sequences are inheritable

Next, we examined the impact of male germline eccDNA on the offspring. To do so, we collected cauda epididymal sperm from AAV-Cre-injected eccDNA mice 4 weeks after AAV injection and performed in vitro fertilization (IVF) using wild type (WT) mouse oocytes (**Figure 2A**). Sperm from empty AAV-injected eccDNA mice (Cre-negative) were used as negative controls. Intriguingly, across two independent experiments, we observed EGFP mRNA in a subset of embryos at the 2-cell stage from the AAV-Cre group (26 out of a combined 67 embryos from the 2 independent experiments) using a STARmap-based RNA fluorescence *in situ* hybridization (FISH) approach (Methods), suggesting the presence of eccDNA^EGFP^ sequences in the embryos (**Figure 2B**). In contrast, no EGFP mRNA signals were detected in the 2-cell embryos from the negative control group (0 out of the combined 56 embryos; AAV-Cre vs controls Fisher’s exact p=2e-6) (**Figure 2B**). Similarly, in a separate batch of experiments (n=2), we detected EGFP mRNA signals in the blastocyst embryos from the AAV-Cre group while observing minimum signals in the negative control group (12 out of the combined 62 embryos in the AAV-Cre group vs. 1 out of the combined 61 embryos in the negative control group; Fisher’s exact p=0.0021) (**Figure S3A**). Interestingly, in both 2-cell and blastocyst embryos from the AAV-Cre group, we did not observe obvious EGFP protein signals because of the low copy number of EGFP mRNA transcripts in the cells (**Figure 2B & S3A**) which might be due to the silencing of the CAG protomer in the eccDNA^EGFP^ sequence.

**Figure 2.**
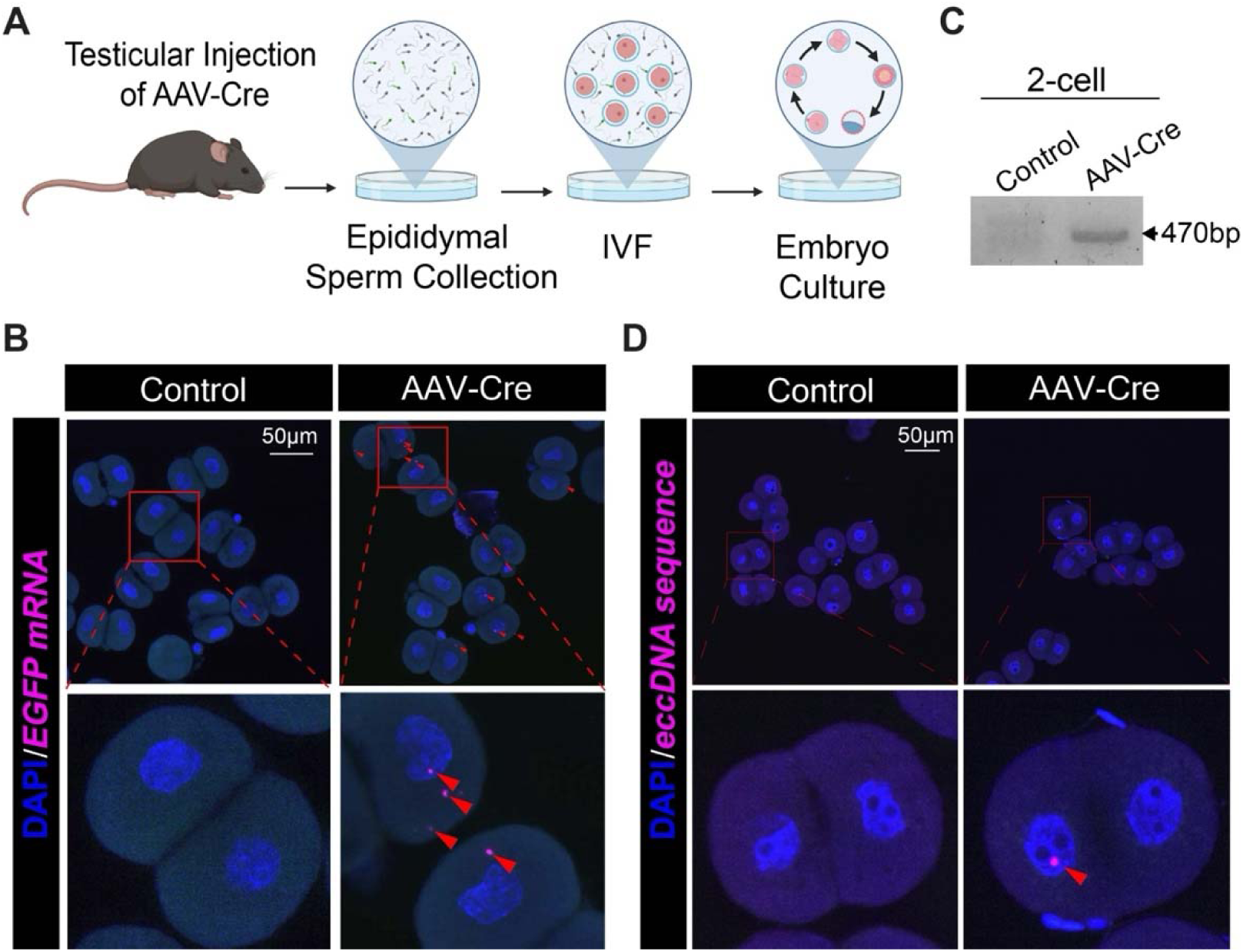
Male germline eccDNA sequences are inheritable. (A) Schematic illustration of the in vitro fertilization (IVF) procedure using sperm from the AAV-Cre injected mice. Image was created using BioRender. (B) Representative RNA FISH images of 2-cell embryos. Red arrow heads denote EGFP mRNA. (C) Positive PCR signals of eccDNA^EGFP^ junction sequence were identified in the DNA of both the 2-cell and blastocyst embryos generated using sperm from the AAV-Cre injected eccDNA mice. (D) Representative RCA-based DNA FISH images of 2-cell embryos. The red arrowhead denotes the eccDNA^EGFP^ sequence.

To further confirm the presence of eccDNA^EGFP^ sequences in the embryos from the AAV-Cre group, the following three orthogonal experiments were performed. First, in another batch of IVF experiments (n=2), we isolated DNA from the 2-cell and blastocyst embryos generated using AAV-Cre-treated sperm and empty AAV-treated sperm, respectively, and performed PCR using primers targeting the junction region of eccDNA^EGFP^. Electrophoresis of the PCR products showed a band with the correct size in the AAV-Cre treated group but not in empty AAV-treated control group (**Figure 2C & S3B**). The sequence identities of the DNA bands were confirmed by Sanger sequencing.

Second, we developed a rolling circle amplification (RCA)-based DNA FISH approach to target the junction region of eccDNA^EGFP^ (Methods). We first tested this approach on HEK293T cells transfected with plasmids containing the eccDNA cassette and the Cre recombinase. Positive signals were observed in cells containing both the eccDNA cassette and the Cre recombinase but not in cells that contained only the eccDNA cassette (**Figure S3C**), demonstrating the specificity of this approach in recognizing the eccDNA^EGFP^ sequence. We then applied this approach to IVF-generated 2-cell embryos. As expected, positive signals were observed in a subset of embryos from the AAV-Cre group (7 out of the 56 embryos) but not in the embryos from the negative control group (0 out of the 61 embryos; AAV-Cre vs controls Fisher’s exact p=0.0047) (**Figure 2D**). The slight difference in the percentage of signal-positive 2-cell embryos between the STARmap-based RNA FISH approach (**Figure 2B**) and the RCA-based DNA approach (**Figure 2D**) may be due to the difference in the sensitivity of these two approaches in detecting the target molecules (EGFP mRNA *vs*. the eccDNA^EGFP^ sequence).

Lastly, to examine if the embryo genome contains the eccDNA^EGFP^ sequence, we performed Nanopore long-read sequencing of the whole embryo genome from 28 blastocysts in the AAV-Cre group. To overcome the limitation of low DNA input, we performed multiple displacement amplification (MDA) of the embryo DNA before Nanopore sequencing (Methods). Analysis of the sequencing data identified several embryo genome insertion events with each event supported by at least two independent chimeric sequences that contain both the eccDNA^EGFP^ sequence and the genomic sequence (**Table S1**). However, since random hexamers in the MDA assay may occasionally mis-prime displaced strands, we could not rule out the possibility that some of the chimeric sequences were MDA artifacts. Nonetheless, our data together suggest that male germline eccDNA sequences can be transmitted to the offspring.

## eccDNA-mediated insertions are common in the human germline genome

Having shown that eccDNA can integrate into the mouse germline genome, we next asked whether eccDNA can leave a heritable sequence footprint in the human genome. To this end, we extracted sequences of polymorphic insertions from the 1000 Genomes Project (1kGP) long-read structural variant call set [19, 20] and aligned them back to the human genome reference (hg38). To identify eccDNA-mediated insertions, we examined the alignments for split sequences with non-colinear mappings, inferred their most likely source intervals, and retained only high-confidence, nonrepetitive, and non-gene duplication events with a unique source sequence (**Figure 3A & 3B**). This screen identified 82 nonredundant putative eccDNA-mediated insertions (**Table S2**) among a total of 327 filtered source-resolved insertions (25.1%) (**Figure 3B**). Two examples of putative eccDNA-mediated insertions identified in the human population are shown in **Figure S4A**.

**Figure 3.**
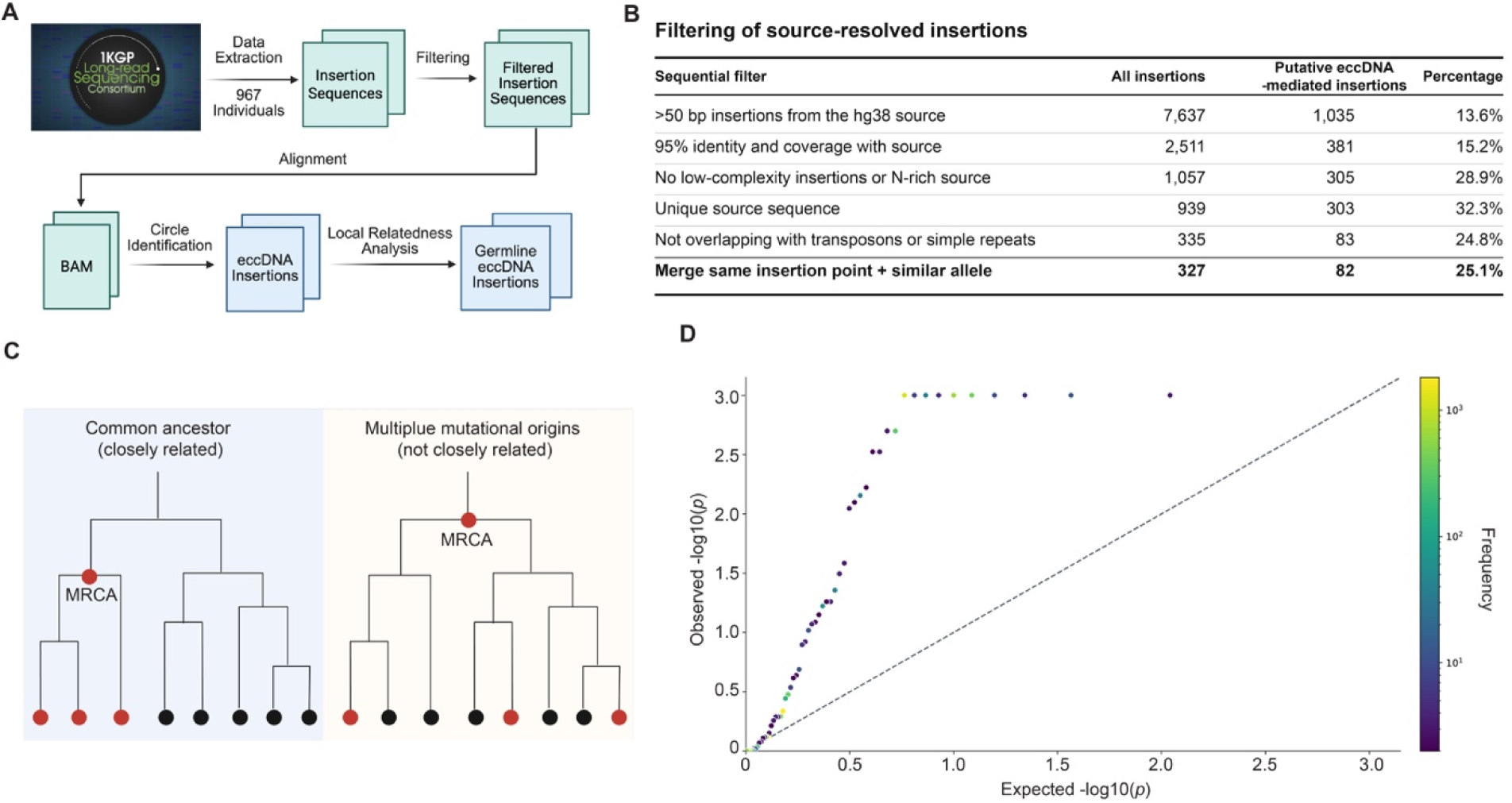
eccDNA-mediated insertions are an important source of human genome structural variants. (A) Workflow to identify germline eccDNA insertions from the 1kGP long-read whole genome sequencing data. (B) Effect of filtering on the fraction of eccDNA-mediated insertions in the total human genome insertions. (C) Plot showing how the number of samples under the MRCA of a set of haplotypes carrying a putative insertion is a measure of the relatedness of the haplotypes in question. Red circles represent haplotypes carrying the structural variant while black dots represent non-carrier haplotypes. Image was created using BioRender. (D) Quantile-quantile plot of empirical p-values for the local MRCA-clade statistic. Expected and observed values are shown on the log p scale. Points above the diagonal indicate putative eccDNA-mediated insertions whose carrier haplotypes have smaller flanking MRCA clades than expected under the matched null model, suggesting greater local relatedness than would be expected if carriers were randomly assigned while preserving allele count, genomic position, and heterozygote/homozygote structure.

Because 55 of the 82 putative eccDNA-mediated insertions were present in more than one individual and passed quality control, we asked whether these alleles were inherited from a common ancestor. We reasoned that if an eccDNA-mediated insertion was transmitted through the germline within a lineage, then the individuals who carry this insertion (carriers) should share the surrounding haplotype more often than expected by chance (i.e., high local relatedness). The local relatedness between a pair of haplotypes can be reflected by their distance in a coalescent tree under a most recent common ancestor (MRCA) (**Figure 3C**). Using an analysis based on the ancestral recombination graph (i.e., a collection of coalescent trees) [21] of single-nucleotide variants flanking the shared insertion sites (Methods), we found that 48 of 55 shared eccDNA-mediated insertions showed excess local sequence sharing between their corresponding carriers (**Figure 3D**), meaning that these 48 eccDNA-mediated insertions were transmitted through the human germline instead of being independent somatic events or recurrent technical artifacts. Together, our analysis shows that eccDNA-mediated insertions is an important source of structural variations in the human germline.

Next, we examined the features of eccDNA-mediated human genome insertions. First, out of the 82 putative eccDNA-mediated insertions, approximately 80% of them are in distal intergenic regions and introns of the human genome while the remaining ones are at the sites of regulatory elements such as the promoters and the 3’UTRs (**Figure S4B**), suggesting the potential impact of eccDNA insertions on human genome functions. Second, the inserted sequences are enriched for terminal microhomology of >=5 bp (allowing one mismatch; 23/82, 28.0% for eccDNA-mediated insertions versus 10/245, 4.1% for filtered non-eccDNA-mediated insertions; Fisher’s exact p=1.3e-08), indicating that at least of a subset of eccDNA were inserted into the linear genome through a microhomology-mediated DNA repair mechanism. Third, for eccDNA-mediated human genome insertions, the origin and the insertion site are often located on the same chromosome (intra-chromosomal insertions) and only 2 out of 82 are inter-chromosomal insertions (**Table S2**). This bias towards intra-chromosomal insertions stands even in comparison to filtered non-eccDNA-mediated insertions (80/82, 97.6% for eccDNA-mediated insertions versus 219/245, 89.4% for filtered non-eccDNA-mediated insertions; Fisher’s exact p=0.022). Lastly, 78 of the 80 intra-chromosomal insertion sites are within 10 Mb from the sources. The inserted sequences are also almost always in the same orientation as the source sequences (79/80, 98.8% for eccDNA-mediated insertions versus 162/219, 74.0% for filtered non-eccDNA-mediated insertions; Fisher’s exact p=8.3e-08). These features suggest that the genomic location of the source of origin may impact the location and orientation of the eccDNA-mediated genome insertion.

## Poly (ADP-ribose) polymerases regulate male germline eccDNA formation

Having established the role of eccDNA in shaping the mammalian germline genome, we next investigated the mechanism of germline eccDNA biogenesis. A common approach to identify molecular factors that may contribute to the generation of germline eccDNA would be to perturb candidate factors (e.g., by knocking down candidate genes) and observe if these perturbations would impact germline eccDNA biogenesis. However, limited perturbation experiments can be done on the human germ cells in vivo due to both experimental and ethical constraints. To overcome these challenges, we looked for naturally occurring perturbations on the human male germ cells. Previous studies have shown that diabetes is coupled with an increased oxidative stress, causing sperm nuclear and mitochondrial DNA damage [22, 23]. Thus, diabetes represents one of naturally occurring perturbations to the human male germline by inducing germline genome instability.

We first set out to quantify sperm eccDNA content from patients diagnosed with diabetes in comparison with that of healthy individuals using a well-established endogenous eccDNA quantification workflow with minor modifications [8, 24]. Briefly, total sperm DNA was subject to a thorough exonuclease treatment to digest the linear DNA. To ensure a complete removal of the linear DNA and a significant enrichment for circular DNA, we used qPCR to monitor the copy number changes of genes on the linear genome and genes on the circular mitochondrial DNA (mtDNA). Only after confirming that the linear DNA signal was no longer detected and that there was at least a 60-fold increase in the enrichment of the circular mtDNA after exonuclease digestion did we continue to the next step. The remaining DNA was tagmented with sequencing adaptors using Tn5 transposases followed by PCR amplification. This amplification approach overcomes the size bias of the RCA-based approach used in the previous circular DNA extraction protocols because smaller DNA circles are preferentially amplified than larger ones by RCA [10]. The amplification products were subject to Illumina short read sequencing and the genome coordinate of each eccDNA was identified using Circle-Map [25]. We applied this approach to 5 sperm samples from 3 healthy adult males and 3 samples from 3 diabetic patients at a similar age. With each sample receiving approximately 200 million sequencing reads, we found that the number of sperm eccDNA was significantly elevated in patients with diabetes when compared with healthy men (**Figure S5A**). Encouraged by this result, we then quantified sperm eccDNA of samples from an additional 4 healthy and 4 diabetic men at a similar age. With approximately 1 billion sequencing reads for each sample, we were able to identify more eccDNA per sample compared to samples sequenced at ∼200 million reads. Consistently, these diabetic patients had significantly more sperm eccDNA than their healthy counterpart (**Figure 4A**). Together, despite patient heterogeneity and small sample sizes, these data suggest that diabetes may lead to the production of germline eccDNA.

**Figure 4.**
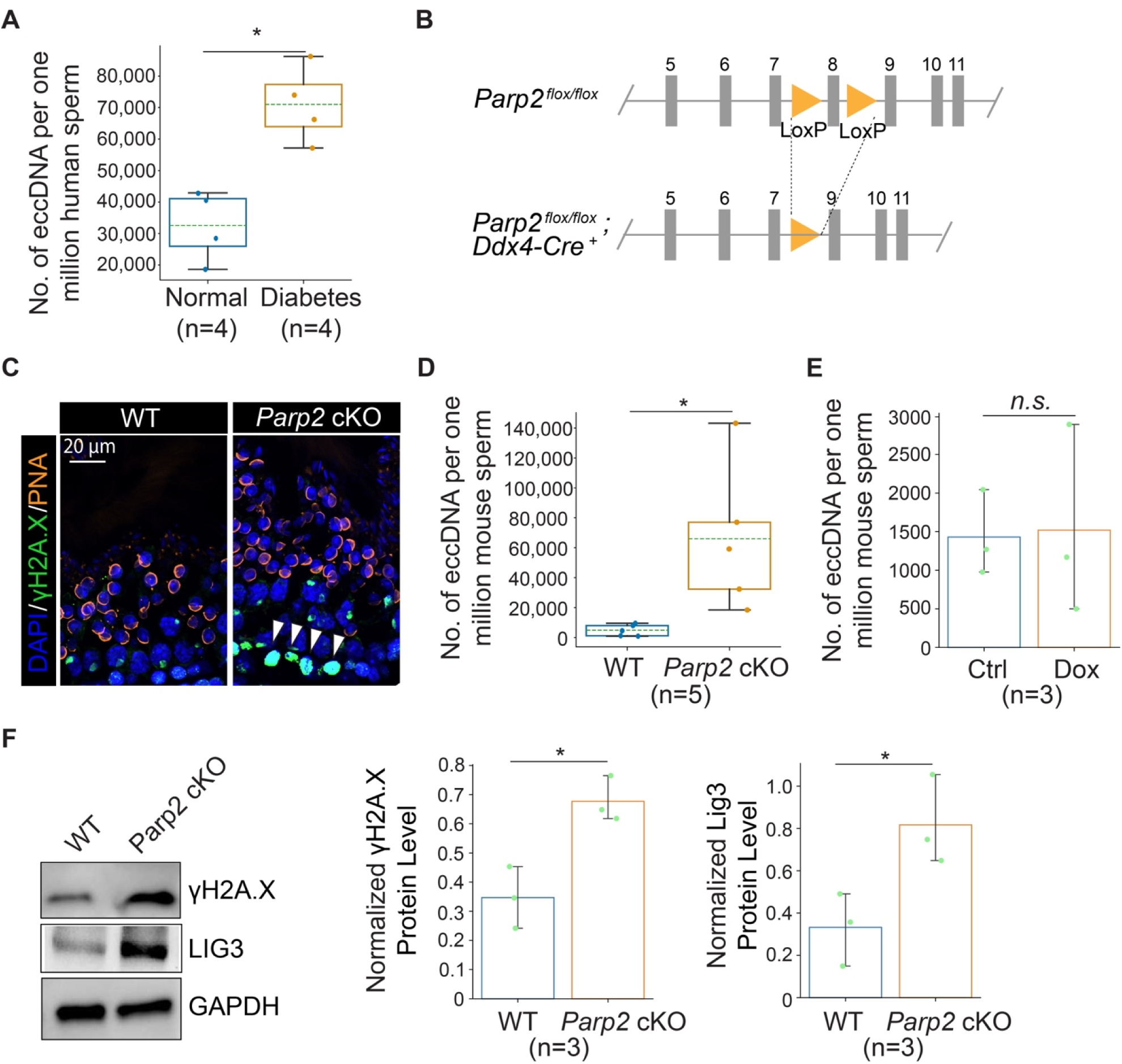
The PARP2 contributes to eccDNA biogenesis in the mammalian male germline. (A) The number of sperm eccDNA in individuals with different health status. *p* values were calculated using a two-tailed Mann-Whitney U tests. *, *p* < 0.05. (B) A germline-specific *Parp2* knockout mouse model was generated by specifically deleting the exon 8 of the *Parp2* gene in the germ cells using *Ddx4* promoter-driven Cre recombinase. (C) Representative images of the seminiferous epithelium of the WT and *Parp2* cKO testis. The γH2A.X signals are in green. White arrow heads denote increased DBS signals in the germ cells of the *Parp2* cKO testis. The acrosomes were visualized with peanut agglutinin (PNA). (D) Bar graph showing the amount of sperm eccDNA in the WT vs. *Parp2* cKO group. The *p* value was calculated using a two-tailed Mann-Whitney U test. *, *p* < 0.05. (E) Bar graph showing the quantification of sperm eccDNA count of control vs. Dox-injected testis. *P* values were calculated using a two-tailed Mann Whitney U test. n.s., not significant. (F) WB analysis of γH2A.X, LIG3, and GAPDH protein levels in testes of WT and *Parp2* cKO mice. *p* values were calculated using a two-tailed Mann Whitney U test. *, *p* < 0.05.

To identify potential molecular players in the diabetes-induced germline eccDNA production, we turned to a previous study [26]. In this study, single cell RNA sequencing (scRNA-seq) of testis samples from diabetic patients was performed and the data was compared with the testis scRNA-seq data of healthy individuals [27] to identify differentially expressed genes (DEGs) in the germ cells at each stage of the development. We screened these DEGs for genes that are known to be involved in genome integrity. Intriguingly, multiple poly (ADP-ribose) polymerase genes (*PARP*s) were present in the DEG list. PARP proteins especially PARP1 and PARP2 were previously reported to show DNA repair activities in the male germ cells when DNA double-strand breaks (DSBs) occur due to oxidative stress, chromatin remodeling, or cell death [28]. Thus, we hypothesized that PARPs may regulate the biogenesis of germline eccDNA. To test this hypothesis, we generated germline-specific *Parp2* knockout mice (*Parp2* cKO) by crossing a *Ddx4*-Cre mouse line with a floxed *Parp2* mouse line (**Figure 4B & S5B**). The reason we chose *Parp2* as our primary target was because the whole body *Parp2* KO male mice were reported to have fertility issues [29]. As expected, *Parp2* cKO mice had elevated DNA damages in the developing male germ cells compared to those of the WT male mice as shown in the γH2A.X (a marker for DSBs) immunofluorescence staining (**Figure 4C)** and TUNEL assay (**Figure S5C**). More importantly, the cauda epididymal sperm of *Parp2* cKO mice contained significantly more eccDNA than the WT sperm (**Figure 4D**, each sample was sequenced at approximately 600 million reads), suggesting that PARP2 may be involved in male germline eccDNA biogenesis.

To examine if the increase in the amount of sperm eccDNA in the *Parp2* cKO mice was specific to *Parp2* KO or was simply due to DNA damages of the germline genomes, we injected doxorubicin (Dox), a DNA damaging agent, to the seminiferous tubules of 3-week-old WT male mice through the rete testis, with the injection of the equal amount of DMSO to a different set of 3-week-old mice as vehicle controls. 4 weeks after injection, we performed TUNEL assay and γH2A.X immunofluorescence staining on the cross-sections of injected testes and quantified the sperm eccDNA content. Interestingly, although elevated DNA damage was observed in the germ cells in the Dox group (**Figure S5D & S5E**), the quantity of sperm eccDNA in the Dox group was at a similar level with that in the control group (**Figure 4E**, each sample was sequenced at approximately 300 million reads). This suggests that damages to the germline genome alone do not simply lead to more germline eccDNA production and that other molecular processes may contribute to *Parp2* KO-induced germline eccDNA biogenesis. Indeed, by performing Western Blot (WB), we observed a significant increase in the protein levels of γH2A.X and DNA ligase 3 (LIG3) in *Parp2* cKO testes compared to the WT testes (**Figure 4F**). In contrast, no increase in the protein level of LIG3 was observed in the Dox-treated testis (**Figure S5F**). LIG3 is a known player in the ligation and circularization of the linear genome fragments to promote eccDNA formation through microhomology-mediated end joining [16, 30, 31]. Together, our data suggest that PARP2 may regulate germline eccDNA biogenesis by influencing genome stability and LIG3 level in the germ cells.

## Discussion

The viability of any species including the human requires that the germline genome is kept stable as it is transmitted across generations by the germ cells. Failure to safeguard the genome integrity and stability would lead to inherited diseases and infertility. Thus, a better understanding of the mechanisms that alter the germline genome is crucial to ensure reproductive health and our continuation as a species.

In this work, we investigated if germline eccDNA can alter the mammalian germline genome. To do this, we established a reporter eccDNA mouse model which allows for the in vivo production of reporter eccDNA with a known sequence identity in a cell type-specific manner. Coupled with long-read sequencing and imaging, this model opens new ways of tracing and tracking the behavior and fate of eccDNA in vivo. We anticipate that this mouse model will serve an invaluable tool to researchers who are interested in studying the biology of eccDNA in various cellular and tissue context. By leveraging this mouse model, we showed that eccDNA generated in the developing male germ cells can re-integrate into different parts of the germline genome and transmit across generations. Through population genetics analysis, we also found that eccDNA-mediated, inheritable germline genome insertions are common in the human population.

Our observation of eccDNA germline integration and transmission echoes previous studies in other species. For example, the phenomenon of ribosomal DNA (rDNA) magnification (i.e., an expanded rDNA copy number) was discovered in the Drosophila male germline over 50 years ago [32]. And the amplification and genome integration of extrachromosomal rDNA circles have been proposed as a potential mechanism to increase chromosomal rDNA copy number [33]. Thus, eccDNA-mediated insertions may represent a conserved mechanism of germline genome alterations across species.

The mechanism through which eccDNA integrate into the linear germline genome is yet to be defined. For an eccDNA to integrate into the linear genome, there must be an accessible entry point in the genome. A pre-existing DSB in the linear chromosome can act as such an entry point [34]. During the mammalian male germ cell development, there are two critical temporal windows during which the male germline intentionally compromises its own genome integrity through programed DSBs: meiotic prophase I [35, 36] and the haploid histone-to-protamine transition [37, 38]. Work is underway to investigate if these programmed DSBs provide entry points for germline eccDNA integration.

By sequencing sperm samples from healthy individuals and diabetic patients and analyzing human testis scRNA-seq datasets, we proposed a link between the PARP family proteins and germline eccDNA biogenesis. KO of *Parp2* in the mouse germline leads to an increase in the protein level of LIG3 and an elevated amount of germline eccDNA. Work is underway to investigate if PARP1 is also involved in eccDNA biogenesis and if changes in PARP protein expression may impact the PARylation of LIG3. It’s worth noting that PARPs may not be the only mechanism of germline eccDNA biogenesis as recent work suggests that the meiotic recombination process may also be associated with germline eccDNA formation [7].

In summary, our work establishes eccDNA as a previously underappreciated mechanism of mammalian germline genome alterations and provides insights into germline eccDNA biogenesis, with important implications for human health and genome evolution.

## Limitations of the study

The synthetic nature of the mouse eccDNA^EGFP^ means that its behavior may not fully reflect that of the endogenous eccDNA. For example, driven by the CAG promoter, the EGFP sequence in eccDNA^EGFP^ can be transcribed in the developing germ cells. A recent work suggests that approximately 2% of human sperm eccDNA contain entire genes and approximately 4% contain promoter regions and an initial exon or exons [7]. It remains to be determined if these human germline eccDNA can transcribe their genetic content. In addition, the sample size of the human sperm eccDNA quantification in the healthy and diabetic individuals was small. A large-scale, systemic investigation on how different disease categories, genetic background, age and other factors may influence sperm eccDNA quantity is needed to fully establish the link between germline eccDNA and patient health.

## Methods Human samples

Adult human sperm samples were obtained from the Fertility & Advanced Reproductive Medicine Clinic at UT Southwestern Medical Center (UTSW). At the clinic, couples who underwent IVF were required to leave semen samples cryopreserved at the clinic in case they were unable to give a fresh specimen on the day of IVF. This has resulted in a large collection of banked backup semen samples from patients with various health status at the clinic. According to standard clinical regulations, these backup semen samples should be discarded after the successful completion of the IVF cycles following patient consent. An IRB-approved protocol allowed us to gain access to these unused backup semen samples for basic research purposes. Adult human testicular samples were obtained through the University of Pittsburgh Health Sciences Tissue Bank and Center for Organ Recovery. All samples were de-identified.

## Animals

All research protocols involving mice were reviewed and approved by the Institutional Animal Care and Use Committee UTSW. The eccDNA mice were generated at Cyagen. *Parp2*^flox/flox^ mice were provided by the Kraus Laboratory at UTSW. *Ddx4-Cre* mice were obtained from the Namekawa Laboratory at the University of California, Davis. Mouse IVF experiments were performed using wild-type B6D2F1 mice (The Jackson Laborator; Cat. #: 100006). All mice were housed under standard pathogen–free conditions (20–22°C, 50–70% humidity, 12-h light/dark cycle) with free access to food and water.

## Genotyping and RT–PCR

Tail biopsies were collected from pups at weaning. Genomic DNA was extracted using the Quick-DNA Miniprep Plus Kit (Zymo Research; Cat. #: D4069). PCR was performed using Q5 Hot Start High-Fidelity 2× Master Mix (NEB; Cat. #: M0494). Total RNA was extracted from the mouse testis, sperm, or embryos by using the AllPrep DNA/RNA Mini Kit (Qiagen; Cat. #: 80204), and cDNA was synthesized using Maxima H Minus Reverse Transcriptase (Thermo Fisher Scientific; Cat. #: EP0753) according to the manufacturer’s instructions. All primer sequences used for genotyping and RT-PCR are listed in **Table S3**.

## Cloning

The eccDNA reporter plasmid used for generating the eccDNA mice were synthesized by Genscript. The AAV-Cre construct was generated using the pX602 plasmid as the backbone. The sequence between the two AAV ITRs was removed using NsiI (NEB; Cat. #: R3127) and NotI (NEB; Cat. #: R3189) restriction enzymes, and was cloned between the ITRs and a CAG promoter–driven Cre cDNA followed by an SV40 poly(A) signal. A WPRE element was inserted between the Cre coding sequence and the SV40 poly(A) signal. The AAV-CRISPR plasmid was was a gift from Yeh-Hsing Lao (Addgene plasmid # 207878). gRNA sequence Cr1 and Cr2 were inserted between the U6 promoter and the SaCas9 guide RNA scaffold following the digestion of the backbone with BbsI (NEB; Cat. #: R3539) and PaqCI (NEB; Cat. #: R0745). The gRNA sequences are as follows: Cr1:5′-CGCCTGTCAGTTAACGGCAGC-3′; Cr2:5′- ATTATTGCTTGTGATCCGCCT-3′

## Cell culture and transfection

Human embryonic kidney HEK293T cells were cultured in Dulbecco’s modified Eagle medium (DMEM; Thermo Fisher Scientific; Cat. #: 11995073) supplemented with 10% fetal bovine serum (FBS; Thermo Fisher Scientific; Cat. #: A5256701) and penicillin/streptomycin (100 U/mL penicillin and 0.1 mg/mL streptomycin) in a humidified incubator at 37°C with 5% CO₂. Cells were typically passaged every 2–3 days at a 1:4 ratio when cultures reached approximately 90% confluence. For passaging, cells were washed twice with calcium- and magnesium-free PBS (1x), detached with 0.05% Trypsin–EDTA for 3–5 min at 37°C, and resuspended in fresh culture medium. For plasmid transfection in a 10-cm dish, 10 μg plasmid DNA was mixed with 1 mL Opti-MEM in one tube. In a separate tube, 40 μL PEI solution (1 mg/mL in water, pH 7.0; Polysciences; Cat. #: 23966) was mixed with 1 mL Opti-MEM. The two solutions were combined, mixed thoroughly, incubated for 10 min at room temperature, and added dropwise to the dish.

## AAV production and purification

AAV9 was produced using AAV-Pro 293T cells (Takara; Cat. #: 632273) cultured in 15-cm dishes. Cells were plated one day before transfection at ∼50% confluence to reach 80–90% confluence on the day of transfection. For each 15-cm dish, 10 μg of AAV-mCherry, AAV-Cre or AAV-CRISPR vector, 10 μg of pAAV2/9 (Addgene; Cat. #: 112865), and 20 μg of pAdDeltaF6 (Addgene; Cat. #: 112867) were mixed with 1 mL Opti-MEM in one tube. In parallel, 160 μL of PEI solution (1 mg/mL in water, pH 7.0) was mixed with 1 mL of Opti-MEM in a second tube. The two solutions were combined, mixed thoroughly, incubated for 10 min at room temperature, and then added dropwise to the dishes. At 96 h post-transfection, cells were scraped and pelleted by centrifugation at 500x g for 10 min. The supernatant was collected and supplemented with one-quarter volume of PEG precipitation solution (40% PEG8000, 2.5 M NaCl), mixed thoroughly, and incubated on ice for 2 h. The PEG/media mixture was centrifuged at 3,000x g for 40 min at 4°C, and the supernatant was discarded. The precipitates were resuspended in 1.5 mL of PBS containing 15 μL of 100 mM PMSF (Thermo Fisher Scientific; Cat. #: 36978), subjected to three freeze–thaw cycles between liquid nitrogen and 37°C, and combined with PEG precipitates. The mixture was treated with 50 μL of DNase I (10 mg/mL) at 37°C for 1 h, followed by centrifugation at 12,000 × g for 1 min at 4°C to obtain clarified viral lysate. AAV particles were purified using a discontinuous iodixanol (Fisher Scientific; Cat. #: NC1059560) gradient. Iodixanol at different concentrations were sequentially loaded into Type 70.1 Ti tubes (1.5 mL of 17% and 25%, and 2 mL of 40% and 60%). Viral particles were carefully layered on top without introducing air bubbles, and gradients were centrifuged at 350,000x g for 2 h 25 min at 4°C (Beckman Coulter; 70.1 Ti rotor). The band at the 40%/60% interface was collected in small fractions and varified by qPCR. Virus-containing fractions were diluted 1:5 in PBS, concentrated using Amicon Ultra-15 100-kDa centrifugal filters at 2,600x g at 4°C, and washed three times with DPBS to a final volume of 50–100 μL. The purified viruses were clarified (20,000x g, 5 min), aliquoted, and stored at −80°C. The final viral titer was determined by qPCR.

## Rete testis microinjection

All procedures were performed under aseptic conditions using sterilized surgical instruments. Mice were anesthetized with inhalation isoflurane. Ophthalmic ointment was applied to prevent corneal drying, and anesthesia depth was confirmed by the absence of pedal reflex. Perioperative analgesia was administered. After shaving and disinfecting the abdominal region, a midline ventral incision (approximately 8-14 mm) was made above the preputial glands, followed by incision of the abdominal musculature. The abdominal fat pad was gently elevated to exteriorize the testes. Under a stereomicroscope, the efferent duct connecting the testis and epididymis was identified. A fine glass microinjection needle was directed toward the rete testis and inserted carefully. For viral delivery, 5 μL of AAV solution (titer at ∼10¹³ GC/mL) containing 1:50 diluted 0.5% Fast Green (Sigma-Aldrich; Cat. #: F7252) was injected to visualize the filling process, ensuring that approximately 70-75% of the seminiferous tubules were infused. Following injection, the testis was returned to the abdominal cavity. The muscle layer was closed using absorbable sutures, and the skin was sealed with wound clips. Mice were allowed to recover individually in sterile cages placed on a 37 °C heating pad and monitored until fully awake. Animals were subsequently housed individually, maintained under clean conditions, and monitored for recovery and wound healing. Mice were maintained for 7-8 weeks post-injection before downstream analyses. For doxorubicin treatment, the drug was injected into the seminiferous tubules of wild-type C57BL/6J mouse testes, with an equal volume of DMSO used as the vehicle control.

## Western blotting

Proteins were extracted from mouse testes using RIPA lysis buffer (Thermo Fisher Scientific; Cat. #: 89900) supplemented with 1% (v/w) protease inhibitor cocktail (Thermo Fisher Scientific; Cat. #: 78429). Protein lysates were separated by SDS–PAGE and transferred onto PVDF membranes, which were blocked with 5% non-fat milk in TBST for 2 h at room temperature, followed by overnight incubation with primary antibodies at 4°C. The following primary antibodies were used: γH2A.X (Thermo Fisher Scientific; Cat. #: MA1-2022; 1:2000 dilution), LIG3 (Abcam; Cat. #: ab313374; 1:2000 dilution), and GAPDH (Thermo Fisher Scientific; 1:5000 dilution). After washing three times with TBST, membranes were incubated with corresponding HRP-conjugated secondary antibodies for 2 hr at room temperature. Signals were developed using ECL Western Blotting Substrate (Bio-Rad; Cat. #: 1705061) and visualized using a Bio-Rad Western Blotting Detection System.

## Immunofluorescence staining

Testes were fixed in 4% paraformaldehyde overnight at 4°C, cryoprotected in 30% sucrose, embedded in OCT, and cryosectioned at 10 μm. Sections were permeabilized with 0.5% Triton X-100 in PBS for 15 min and blocked with 5% bovine serum albumin (BSA) in PBST for 12 h at 4 °C. Sections were incubated with anti-γH2AX primary antibody (Thermo Fisher Scientific; Cat. #: MA1-2022; 1:200 dilution) overnight at 4°C. After washing with PBST, sections were incubated for 2 h at room temperature with Alexa Fluor 488-conjugated goat anti-rabbit IgG secondary antibody (Thermo Fisher Scientific; Cat. # A-11008) together with PNA (Thermo Fisher Scientific, Cat. #L32459). Nuclei were counterstained with DAPI, and fluorescence images were acquired using a Nikon spinning disk confocal microscope at 40x or 60x magnification.

## TUNEL assay

Mouse testes were fixed in 4% paraformaldehyde for 12 hr, cryoprotected in 30% sucrose at 4°C for 24 hr, embedded in the OCT compound (Sakura; Cat. #: 4583) and stored at −80°C. TUNEL staining was performed on the testis cross-sections using the TUNEL BrightGreen Apoptosis Detection Kit (Vazyme; Cat. #: A112) according to the manufacturer’s instructions. Images were acquired using a Nikon spinning disk confocal microscope at 40x or 60x magnification and analyzed using ImageJ software (v1.50).

## IVF

Female B6D2F1/J mice (6 weeks old; The Jackson Laboratory; Cat. #: 100006) were super- ovulated by intraperitoneal injection of 5 IU of pregnant mare serum gonadotropin (PMSG; Fisher Scientific; Cat. #: NC1663485), followed 48 hr later by 5 IU of human chorionic gonadotropin (hCG; Sigma-Aldrich; Cat. #: C1063). Cumulus–oocyte complexes were collected from the oviducts 12-14 hr after hCG administration and placed into 100-μL droplets of human tubal fluid (HTF; Sigma-Aldrich; Cat. #: MR-070-D) supplemented with 5 mg/mL BSA (Sigma-Aldrich; Cat. #: A3803). Epididymal spermatozoa were obtained from the eccDNA mice whose testes were injected with AAV-Cre and were capacitated in 0.5 mL of fertilization medium (HTF supplemented with 4 mg/mL BSA) at 37 °C for 1 hr. A 10-μL aliquot of capacitated sperm suspension was then added to each fertilization drop containing oocytes to achieve a final sperm concentration of approximately 2x10 /mL. Oocytes retrieved from both oviducts were pooled into a single fertilization drop, and were co-incubated with the sperm for 4 hr. Presumptive zygotes were washed three times in 20-μL droplets of KSOM Mouse Embryo Medium (Sigma-Aldrich; Cat. #: MR-106-D) and subsequently cultured at 37 °C in a humidified atmosphere of 5% CO₂ and 5% O₂.

## Sperm eccDNA purification

Plasmid Safe ATP-dependent DNase kit (BioSearch Technologies; Cat. #: IDE3110K) was used to digest sperm chromosomal linear DNA . For 2 μg of total sperm DNA, 30 units (3 μL) of exonuclease, 4 μL of ATP (25 mM), and 10 μL of 10x reaction buffer were added to reach a reaction volume of 100 μL. The digestion was performed at 37°C for at least 3 days. Each day an additional 4 μL of ATP (25 mM), 0.6 μL of 10x reaction buffer, and 3 μL of exonuclease were added into the reaction mix to continue the enzymatic digestion reaction. The exonuclease treated sample was used to confirm elimination of chromosomal linear DNA and the enrichment of circular DNA by quantitative polymerase chain reaction (qPCR), using chromosomal marker *COX5B* and mtDNA marker *mt-ND2*. qPCR primer sequences are listed in **Table S3**. Each 20 μL of qPCR reaction contains 2 μL of exonuclease-treated sample, primers at a final concentration of 150 nM, 0.2 μL of 100x SYBR Green dye (ThermoFisher Scientific; Cat. #: S7563), and 10 μL of Q5 Hot Start High-Fidelity 2X Master Mix (NEB; Cat. #: S7563). 1 ng of undigested total sperm DNA from the same sample was used as control. The following qPCR reaction condition was used: 30 sec at 98°C, followed by 35 cycles of 10 sec at 98°C, 15 sec at 65°C, and 30 sec at 72°C. If qPCR result indicated the existence of remaining undigested linear DNA, an additional day of exonuclease digestion was performed. If the result confirmed the elimination of linear DNA and the enrichment of mtDNA, the exonuclease in the digestion solution was heat inactivated by incubating the solution at 70 °C for 30 minutes. The digested DNA was purified using a NucleoSpin Gel and PCR Clean-Up Kit (Takara; Cat. #: 740609) and then quantified using a Qubit dsDNA Quantification Assay Kit (ThermoFisher Scientific; Cat. #: Q32851). A Nextera XT Library Prep Kit (Illumina; Cat. #: FC-131-1024) was used to generate a sequencing library following the manufacturer’s instructions except that DNA at 0.4 ng/μL instead of 0.2 ng/μL was used as input. The concentration and size distribution of the sequencing libraries were quantified using the Agilent D1000 High Sensitivity Screentape Kit (Agilent Technologies; Cat. #: 5067-5584) kit on an Agilent TapeStation 2200 system. Libraries were sequenced on an Illumina NextSeq 2000 system or an Illumina NovaSeq X Plus system with a sequencing run cycle of 150 bp/8 bp/8 bp/150 bp (read 1/index 1/index 2/read 2).

## Identification of eccDNA with Circle-Map

The adaptor sequences of the paired end reads were trimmed using BBDuk under BBMap v38.46. The trimmed reads were aligned to the hg38 version of the human genome using BWA MEM v.0.7.5. All downstream BAM/SAM file processing analyses were performed using Samtools v1.6. Circle-Map v.1.1.4 was used to detect eccDNAs by first extracting all soft-clipped, hard-clipped, and discordant reads to a new BAM file using the ‘ReadExtractor’ command and then executing the ‘Realignment’ module while keeping all parameters at their default settings. Mitochondrial DNA was removed from downstream analysis. For the mouse data, a custom realign script was used to identify eccDNA using the output from Circle-Map ReadExtractor as the input data as we have observed a prolonged runtime when using the Circle-Map ‘Realignment’ module.

## Library preparation for Nanopore sequencing

gDNA were extracted by incubating sperm or embryo samples in 200 μL of solid tissue buffer (ZYMO Research; Cat. #: D4069) containing 10 μL of Proteinase K (Thermo Scientific; Cat. #: EO0491) and 10 μL of 0.5 M DTT (Thermo Fisher Scientific; Cat. #: R0861), and incubated overnight at 55°C. Following lysis, an equal volume of phenol:chloroform:isoamyl alcohol (Thermo Fisher Scientific; Cat. #: 15593031) was added, mixed thoroughly, and centrifuged at 13,000x g for 10 min at 4°C. The aqueous phase was transferred to a new tube, incubated with 500 μL of isopropanol at room temperature for 10 min, and centrifuged again at 13,000x g for 10 min at 4°C. The resulting DNA pellet was washed with 1 mL of 70% ethanol, air dried, and resuspended in 50 μL of elution buffer (ZYMO Research; Cat. #: D4069). DNA concentration was quantified using Qubit. For the whole genome long-read sequencing, genomic DNA was fragmented using a g-TUBE (Covaris; Cat. #520079) according to the manufacturer’s instructions. The fragmented DNA was purified using 0.8x AMPure XP beads (Beckman Coulter; Cat. #A63881), yielding DNA fragments with a peak size of approximately 10 kb. For the targeted long-read sequencing, genomic DNA was fragmented using Tn5 transposase. Briefly, partially double-stranded adaptors A and B were prepared by annealing 10 mM Tn5-A (5’-TCGTCGGCAGCGTCAGATGTGTATAAGAGACAG-3’) or Tn5-B (5’-GTCTCGTGGGCTCGGAGATGTGTATAAGAGACAG-3’) oligonucleotides with equal molar amounts of mosaic end oligonucleotides (5’-/5phos/CTGTCTCTTATACA/3ddC/-3’) in annealing buffer (10 mM Tris–HCl pH 7.5, 10 mM NaCl). Annealing was performed at 94°C for 5 min followed by gradual cooling to 10°C at 1°C/min. Tn5 transposases were loaded with an equimolar mixture of adaptors A and B in IDB buffer (10% glycerol, 10 mM Tris-HCl pH 7.5, 20 mM NaCl, 20 μM EDTA, 0.2 mM DTT, 0.02% NP-40) for 30 min at room temperature and stored at −20°C until use. Loaded Tn5 were then used to fragment 1 μg of genomic DNA by co-incubation at 55°C for 10 min. Fragmented DNA was purified using 0.8x AMPure XP beads (Beckman Coulter; Cat. #: A63881), and fragment size distribution was assessed using TapeStation, ensuring that the majority of fragments ranged between 1-5 kb. Libraries were amplified using primers eccDNA-P5 (5’-GAGCTTTGCTAACGGTCG-3’) and eccDNA-EGFP (5’- ATGGCGGACTTGAAGAAG-3’) to selectively enrich for DNA fragments containing the eccDNA^EGFP^ sequences. PCR products were then purified using 0.8x AMPure XP beads, and fragment size and concentration were determined using TapeStation. For embryo whole genome sequencing, multiple displacement amplification (MDA) was performed prior to library preparation due to the limited amount of genomic DNA in embryos. MDA was performed using the REPLI-g Single Cell Kit (QIAGEN; Cat. #: 150343) according to the manufacturer’s instructions. Briefly, genomic DNA was denatured at 65°C for 10 min and neutralized with the stop solution. MDA was then carried out in a 50 µl reaction at 30°C for 2 h, followed by enzyme inactivation at 65°C for 3 min. The amplified DNA was purified using AMPure XP beads and quantified using a Qubit fluorometer. Subsequently, 1.5 µg of amplified DNA was treated with T7 Endonuclease I (NEB; Cat. #: M0302) at 37°C for 60 min to resolve hyperbranched amplification products. The digested DNA was purified using AMPure XP beads before downstream library preparation. Nanopore sequencing libraries were prepared using the Ligation Sequencing Kit (Oxford Nanopore Technologies; Cat. #: SQK-LSK114) according to the manufacturer’s instructions and sequenced on a MinION or PromethION flow cell.

## Nanopore sequencing data analysis

A hybrid reference genome was generated by appending the eccDNA^EGFP^ sequence to the mouse GRCm39 reference genome. Nanopore long reads were aligned to the hybrid reference using minimap2 (v2.28) with parameters optimized for Oxford Nanopore sequencing (map-ont), retaining supplementary alignments and soft-clipped bases. Chimeric reads spanning both eccDNA and genomic sequences were identified using a custom script by aggregating primary and supplementary alignments for each read. After quality filtering (MAPQ ≥ 20 and aligned segment length ≥ 100 bp), eccDNA integration sites on the genome were inferred at single-base based on the relative positions of the eccDNA^EGFP^ sequence and the associated genomic sequence within each read. To account for breakpoint jitters inherent to Nanopore sequencing, breakpoints within 10 bp were merged into a single integration site. An integration site was considered valid only if it was supported by at least two independent sequencing reads. Nonredundant integration sites were annotated using ChIPseeker (v1.32.1) [39] with gene models derived from Ensembl GRCm39 or hg38, defining promoter regions as ±3 kb from transcription start sites (TSS). Genome-wide distributions of integration sites were visualized using circlize (v0.4.17) [40].

## STARmap- and RIBOmap-based RNA detection

The STARmap [41] protocol was adopted to detect EGFP transcripts in the embryos and the RIBOmap [42] protocol was adopted to detect ribosome-associated EGFP transcripts in the mouse testis samples. Probe sequences are listed in **Table S3**. All probes were synthesized by Integrated DNA Technologies (IDT).

Mouse testes were fixed in 4% PFA for 12 hr and cryoprotected in 30% sucrose at 4°C for 24 hr, embedded in OCT, and stored at −80°C. Cryosections (10 μm) were prepared at −20°C using a Leica CM1950 cryostat and mounted onto glass-bottom 24-well plates (Cellvis, P24-1.5H-N)) pretreated with 3-(Trimethoxysilyl) propyl methacrylate (Sigma-Aldrich; Cat. #: M6514) and poly-D-lysine (Sigma-Aldrich; Cat. #: A-003-M). Sections were fixed in 4% PFA in PBS for 15 min at room temperature, permeabilized in cold methanol at −20°C for 1 hr, equilibrated for 5 min at room temperature, and washed twice with PBSTR [PBS containing 0.1 U/μL RNase inhibitor (NEB; Cat. #: M0314L) and 0.1% Tween-20] for 10 min each.

Embryos were fixed in 4% PFA for 30 min at room temperature, washed three times with PBS, permeabilized in cold methanol at −20°C for 1 h, equilibrated at room temperature for 5 min, and washed twice with PBSTR for 10 min each prior to downstream STARmap processing.

Samples were then incubated with 200 μL of 1× hybridization buffer [2× SSC (Thermo Fisher Scientific; Cat. #: 15557044), 10% formamide, 20 mM ribonucleoside vanadyl complex (NEB; Cat. #: S1402S), 0.1 mg/mL yeast tRNA (Thermo Fisher Scientific; Cat. #: AM7119), 0.5% RNase inhibitor, 0.1% Tween-20, pooled padlock probes at 1 nM, and primer oligos at 1 nM]. For RIBOmap, splint probes at 100 nM were also added into the hybridization buffer. Hybridization was carried out at 40°C for 12 hr in a humidified chamber. Samples were washed twice with 300 μL of PBSTR and once with 300 μL of high-salt wash buffer (4× SSC in PBSTR) for 20 min each at 37°C, followed by a final rinse in PBSTR at room temperature. Samples were then incubated with 200 μL of ligation buffer (0.25 U/μL T4 DNA ligase, 0.5 mg/mL BSA, and 0.4 U/μL RNase inhibitor in 1× T4 ligase buffer) at room temperature for 2 hr with gentle shaking. After washing twice with PBSTR, RCA was performed using 200 μL of RCA mix [0.5 U/μL Phi29 DNA polymerase (Thermo Fisher Scientific; Cat. #: EP0094), 250 μM dNTPs (Thermo Fisher Scientific; Cat. #: 18427089), 0.5 mg/mL BSA, and 0.4 U/μL RNase inhibitor in 1× Phi29 buffer] at 4°C for 30 min followed by 30°C for 2 hr with gentle shaking. Samples were washed twice with PBST, then washed three times with wash-10 buffer (10% formamide in 2x SSC) for 10 min each. Amplicons were detected by incubating samples with detection probe solution (100 nM in wash-10 buffer) for 1 hr at 37°C, followed by washing in wash-10 buffer and PBS at 37°C for 20 min each. Nuclei were counterstained with DAPI for 15 min and washed in PBS for 10 min. Images were acquired using a Nikon spinning disk confocal microscope.

## Chromogenic DNA ISH

Probes specifically targeting the junction site of the eccDNA^EGFP^ was designed and synthesized by ACDbio. DNA FISH was performed using the BaseScope Intro Pack Reagent Kit v2 (ACDbio; Cat. #: 323971) according to the manufacturer’s instructions. Briefly, tissue sections were dehydrated through an ethanol gradient (50%, 70%, and 100%) and air-dried. Sections were treated with hydrogen peroxide for 10 min at room temperature, washed in water, and subjected to target retrieval in Target Retrieval Solution at 98-102 °C for 5 min. After washing and dehydration in 100% ethanol, sections were air-dried and incubated with protease III at 40 °C in a humidified chamber for 30 min. Samples were washed and hybridized with the target probes at 40 °C for 2 h. Signal amplification was then performed using sequential hybridization with Amp 1–Amp 8 reagents from the BaseScope v2 amplification system. Signals were developed using RED detection solution for 10 min, followed by water wash and nuclear counterstaining with 50% hematoxylin.

For sperm samples, the epididymis was dissected, and spermatozoa were allowed to swim out into PBS. Sperm were pelleted (3,000 rpm, 6 min), resuspended in 50 μL of PBS, and diluted 1:4 in water. 5-μL drops of sperm were spread onto glass slides and air-dried. Slides were incubated with 100 μL freshly prepared de-condensation buffer (25 mM DTT, 0.2% Triton X-100, and 200 IU/mL heparin in PBS) for 15-18 min in a humidified chamber. De-condensation progress was monitored by phase-contrast microscopy. Processing was stopped when most nuclei appeared dull gray and approximately doubled in nuclear area. Slides were then fixed in 4% PFA (PBS, pH 7) for 15 min, washed in PBS, and air-dried prior to DNA ISH processing.

## RCA-based DNA FISH

RCA-based DNA FISH was performed on HEK293T cells and IVF embryos. Samples were fixed with 4% paraformaldehyde, permeabilized with 0.5% Triton X-100, and dehydrated through a graded ethanol series. To generate single-stranded DNA at the target locus, samples were sequentially incubated with Nb.BssSI (NEB, Cat. #: R0681) (2 μl in a 200 μl reaction, 37°C for 60 min) and Exonuclease III (NEB, Cat. #: M0206) (4 μl in a 200 μl reaction, 37°C for 60 min). Padlock and primer probes were hybridized overnight (12–16 hr) at 37°C in hybridization buffer containing 20% formamide, 2× SSC, 0.1% Tween-20, and 0.5 mg/ml BSA, followed by ligation with T4 DNA ligase (NEB, Cat. #: M0202) in a 200 μl reaction for 2 h at room temperature. RCA was carried out in a 200-μl reaction containing 1x Phi29 buffer, 0.5 U/μl Phi29 DNA polymerase, 250 μM dNTPs, and 0.5 mg/ml BSA, with pre-incubation at 4°C for 15 min followed by amplification at 30°C for 4 h. RCA products were detected by hybridization with a fluorescent detection probe at 100 nM for 1 h at 37°C, followed by DAPI counterstaining and fluorescence microscopy.

## Identification of eccDNA-mediated genome insertions in the human population

Polymorphic insertion alleles were analyzed from the 1kGP long-read structural variant call set [19, 20]. Inserted allele sequences more than 50 bp were extracted and aligned to the hg38 reference genome to infer their most likely source sequence. For each insertion allele, 50-nt perfect-match seeds were sampled every 25 nt from both the reported inserted sequence and its reverse complement. Seed hits to each reference chromosome were extended into exact alignment blocks, and compact block groups on the same reference chromosome and strand were retained when the candidate source span was no greater than 5 kb. Candidate insertions were classified by comparing the order of aligned blocks along the inserted allele with the order of the same blocks along the inferred source interval. Insertions in which the query block order represented a non-zero cyclic shift of the source order were classified as AB->BA, consistent with insertion of a circularly permuted source molecule. Colinear source-resolved insertions that did not show this cyclic block order were retained as the matched control group of non-AB->BA insertions.

The same sequence and reference filters were applied to AB->BA candidates and to the matched denominator. We required at least 95% query coverage and 95% source coverage, with no more than 5% overlap among aligned blocks in either the query or source coordinate system. Low-complexity alleles were removed if the inserted sequence contained a homopolymer tract of 20 bp or longer or had low complexity defined as Shannon sequence entropy below 1.6. Source intervals were also excluded when the corresponding hg38 sequence, assessed across the source interval plus 50 kb of flanking sequence on each side, contained more than 5% N bases. To avoid ambiguous source assignments, source uniqueness was evaluated at the sequence level after quality filtering: candidate insertions were retained only when they were matched to a single unique source sequence. The only retained insertion when the same sequence occurred at more than one genomic locus is orthologous loci between X and Y chromosomes in pseudoautosomal region. We also removed insertions annotated in the variant call set as overlapping transposable elements, simple repeats, variable number tandem repeats, or nested repeat breakpoints. Finally, candidate insertions were collapsed by insertion coordinate and inferred source-sequence identity, and highly similar inserted alleles at the same insertion point were merged. This procedure yielded 327 filtered source-resolved insertion groups, of which 82 contained an AB->BA pattern which we deemed as eccDNA-mediated genome insertions. The remaining 245 formed the filtered non-AB->BA comparison set (i.e., non-eccDNA-mediated genome insertions).

A secondary confidence review was performed for the 82 eccDNA-mediated insertion groups. Ten groups were flagged as lower-confidence cases in which a short segment supports the AB->BA pattern. This category included two groups whose circular-permutation classification was not stable in the reverse-complement control described below, and eight additional groups in which the AB->BA pattern was supported by a small terminal block, overlapping alignment blocks, or another near-boundary block configuration. These 10 groups were retained in the primary count because they passed the same source, repeat, uniqueness, and merging filters as the other AB->BA insertions. However, they were tracked separately as a lower-confidence category.

To test robustness of our approach, we repeated the classification and downstream analyses after replacing each inserted allele sequence with its reverse complement while keeping the same variant identifiers, insertion coordinates, and genotype metadata. Because the source search indexed both orientations of every allele, this control was implemented equivalently by reflecting the query coordinates in the existing alignment blocks and flipping the query strand, followed by rerunning the AB->BA classifier, matched filters, merging, and downstream feature analyses. The control recovered the same final set of 327 insertion/source clusters, with nearly identical AB->BA counts [83 (80 being identical) in the reverse-complement control versus 82 in the reported orientation].

## Estimation of local relatedness around eccDNA-mediated insertions

To investigate whether the putative eccDNA-mediated genome insertions we observed in the 1kGP represent somatic mutations in the blood, sequencing artifacts, or germline mutations, we measured the degree of relatedness of the haplotypes carrying the purported insertions. The logic behind this approach is that if the insertions were inherited from a common ancestor through germline transmission, then we would expect the carrier haplotypes to be closely related. In contrast, we would expect the sequencing artifacts/somatic mutations to be distributed across unrelated haplotypes. For this analysis, we used the 2022 release of the 1000 Genomes Project and subset the sample such that it contains haplotypes only from the individuals present in the 2015 release [43, 44].

Haplotypes in contemporary population cohort represent imperfect copies of haplotypes in the past. As we go further back in time a pair of sampled haplotypes will at some point have the same ancestor, also called the most recent common ancestor (MRCA) of the pair. We say that the haplotype lineages coalesce at the MRCA, thus producing a coalescent tree as shown in **Figure 3C**. Coalescent trees allow us to quantify how closely related a pair of haplotypes are by measuring how quickly they coalesce: a pair of closely related haplotypes will coalesce more recently than a pair of distantly related haplotypes. The same is true when we study the relatedness of more than two haplotypes. An effective metric that captures the degree of relatedness is the number of samples under the most recent common ancestor for the set of haplotypes [45, 46] (also see **Figure 3C**). As the number of samples under the MRCA increases, the relatedness decreases. It’s worth noting that our proposed procedure contains noises which dilute the signal of relatedness. However, this has the advantage of making our test conservative.

In the human population, the haplotypes get reshuffled by meiotic recombination. Thus, different genomic locations have different coalescent trees, and the haplotypes are related by a complex and collection of coalescent trees called an ancestral recombination graph (ARG) [47]. As a result, a pair of homologous chromosomes can have different degrees of relatedness depending on where in the genome we measure the relatedness. We used *tsinfer* to construct the ARG from the 1kGP VCF files and selected point mutations, excluding rare variants (with frequency *k* ≤ 20), in the construction of the tree sequence [21, 43].

Let an observed putative insertion be indexed by *i*, located on chromosome *c_i_*, genomic position *x_i_*, allele count *k_i_*, heterozygous carrier set *H_i_*, and homozygous carrier set *G_i_*. We exclude singletons (*k_t_* = l) from the analysis because local relatedness is undefined for them. If we are unable to reconstruct the ARG in the vicinity of the putative insertion, then we exclude that insertion from our analysis. We also exclude insertions located on sex chromosomes given the difficulty in constructing and analyzing the ARG. Finally, missing haplotypes are excluded from the analysis. From the original 82 putative eccDNA-mediated insertions we are left with 55 (**Figure S6A**). Since humans have two copies of each allele, heterozygote carrier can have the insertion located on either copy of chromosomes *c_i_*. The process of attributing the insertion to a specific chromosome is called phasing. We assume that the phase for the insertions is uncertain, so for a heterozygous carrier, the insertion could lie on either of its two haplotypes. To account for this, we introduce the minimal phase-agnostic MRCA statistic which accounts for this ambiguity by considering all phase-compatible choices and taking the smallest clade that can explain the sampled carrier haplotype. Suppose that *m* individuals carry the same insertion of interest. Each individual *j* ∈ {1,2, *m*} has two haplotypes *h_j,0_* and *h_j,1_*. A heterozygous carrier contributes one of its haplotypes, while a homozygous carrier contributes both of its haplotypes to the sample. Thus, for homozygotes there is no phase ambiguity, and the phase correction only applies to heterozygotes. For one local tree *T*, define a phase-compatible haplotype choice as:

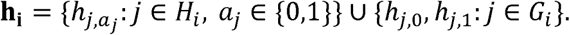

For each choice, let *n_T_*(**h_i_**) be the MRCA node in the local tree of the haplotype choice **h_i_**. The number of sample haplotypes descended from node *n_T_* in tree *T* is denotes as *L_T_*(*n_T_*(**h_i_**)) and the minimum phase-agnostic clade size for that local tree is

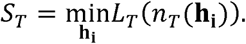

To account for noises in the ARG reconstruction (for example: missed recombination events), which are likely to increase the number of sample haplotypes under the MRCA of **h**_i_, we scan the flanking trees near the putative insertion and select the smallest value of *S_T_* over all possible trees *T* in the flanking interval. In addition, we exclude the central region around the insertion, and any tree interval that intersects it, to account for any possible sequencing errors. As a result, for a focal position *x_i_* the eligible flanking intervals are

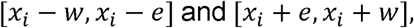

where we selected 2*w* = 150 kb for the width of the flanking interval and 2*e* = 400 bp for the width of the excluded central region (see **Figure S6B**). Let *F_i_* be the set of eligible flanking trees. The relatedness statistic is

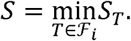

A small value of *S* means that carrier haplotypes can be placed within a small local genealogical clade after accounting for unphased heterozygotes. The raw values of *S* are difficult to interpret, so we want to investigate if the minimal clade sizes are smaller than we would expect by assigning the insertions to random haplotypes while preserving the features of the putative insertions (the observed position *x_i_*, the frequency class *k_i_*, and the observed heterozygote/homozygote structure).

For each putative insertion, we sample a matching set of individuals a total of *N* = 1000 times. The empirical null statistic for the *n^th^* control set is:

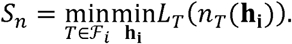

The quantity of interest is the lower-tail empirical p-value, which for the Z-th insertion is:

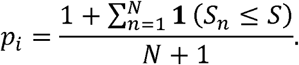

The plus-one correction prevents a *p*-value of exactly zero when no null replicate is as small as the observed statistic. A small p-value means that matching control set has lower relatedness than the carrier set of the putative insertions. In the context of assessing whether calls might be somatic mutations or sequencing artifacts, a small p-value argues against a simple model in which carriers are randomly scattered across the tree. We show the quantile-quantile plot of the p-values in **Figure 3D** and we observe that 48 of 55 putative eccDNA-mediated insertions, or 87.3%, fall above the quantile-quantile diagonal, indicating an excess of small *p*-values relative to the uniform null expectation. If an insertion has multiple mutational origins, then our metric *S* would be unable to differentiate between sequencing errors/somatic mutations and germline mutations. However, given that we see tight clusters for rare putative insertions, we can conclude that they represent true germline mutations inherited from a common ancestor.

## Data availability

The raw sequencing data supporting the findings of this study are available in the NCBI BioProject database with BioProject ID PRJNA1117395.

## Code availability

Custom code is available at https://github.com/HaiqiChenLab/human_sperm_eccDNA and https://github.com/alicodendrochit-alt/eccdna-abba-population-insertions.

## Acknowledgments

We thank Sihan Wu for helpful discussions, Dorothy Mundy for providing assistant with SIM, and Lei Wang and Qianlan Xu for assistant with Nanopore sequencing. H.C. acknowledges support from the Cecil H. and Ida Green Center for Reproductive Biology Sciences Endowment.

## Author contributions

H.C. conceived and supervised the project; X.Z. performed experiments with assistance from M.E., S.R. and A.H.; H.C. and X.Z performed data analysis together with M.E., S.R., R.S., V.S.,

Y.Z. and L.X.; K.S. and O.B. provided human semen samples. K.S., L.X., O.B. V.S. and K.E.O provided consultations. H.C. and X.Z. wrote the manuscript with input from all authors.

---

**Figure S1.**
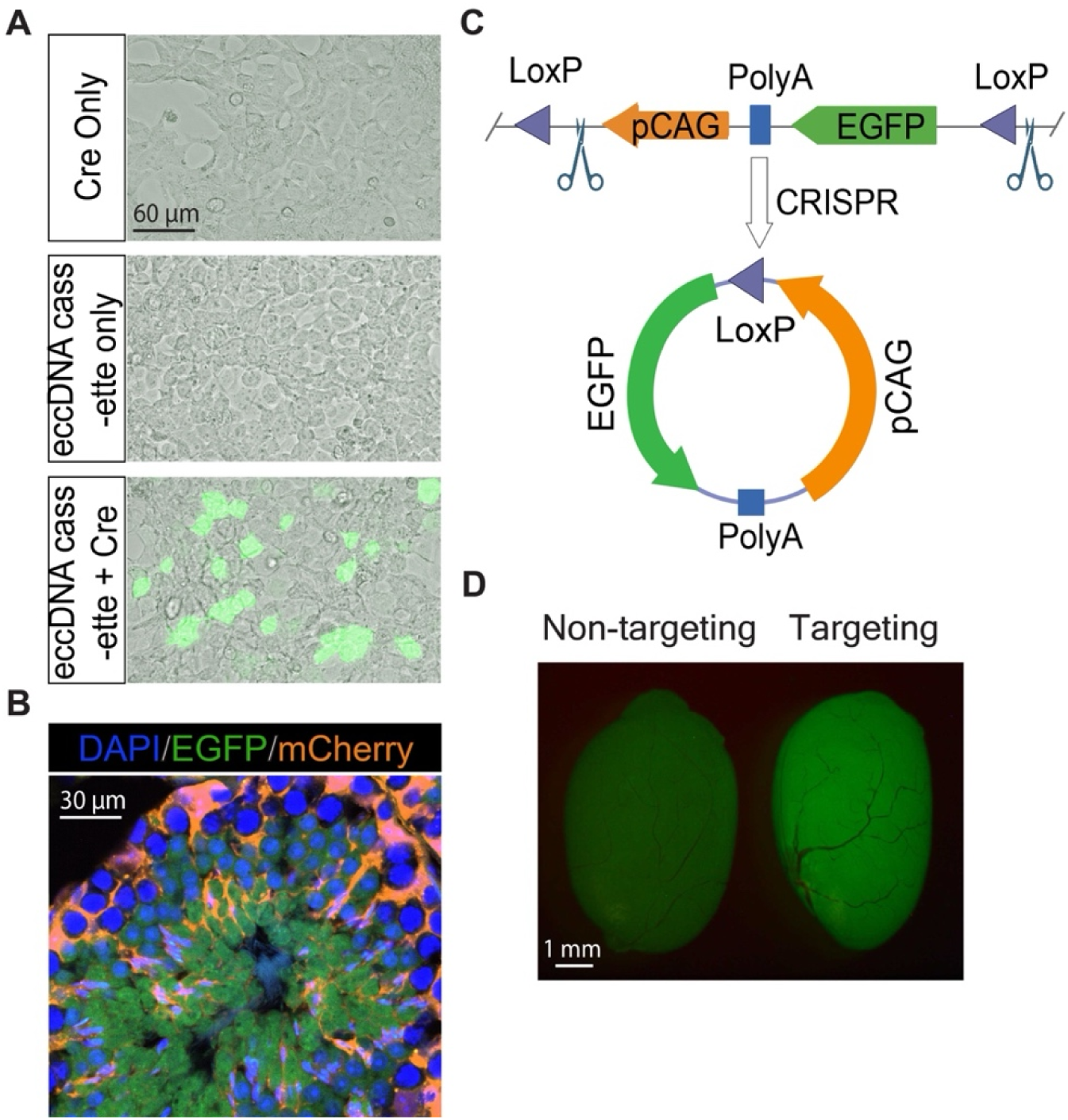
Strategies to induce eccDNA^EGFP^ formation in vivo. (A) Representative images of HEK293T cells transfected with plasmids expressing different components of the eccDNA^EGFP^ system. The green fluorescence indicates the successful generation of eccDNA^EGFP^. (B) Representative image of a seminiferous tubule from a mouse testis injected with AAVs that express mCherry. This experiment demonstrated the feasibility of the rete testis injection method to introduce AAVs to developing germ cells at different developmental stages. Germ cells are genetically labeled with EGFP. As indicated by the mCherry fluorescence, AAVs introduced through the rete testis injection method can infect both Sertoli cells as well as developing germ cells at various developmental stages. (C) Schematic diagram of CRISPR-C mediated eccDNA^EGFP^ formation. (D) Injection of AAVs expressing the SaCas9 protein and eccDNA cassette-targeting sgRNAs to the testis induced eccDNA^EGFP^ formation in vivo as indicated by the green fluorescence signals in the injected testis.

**Figure S2.**
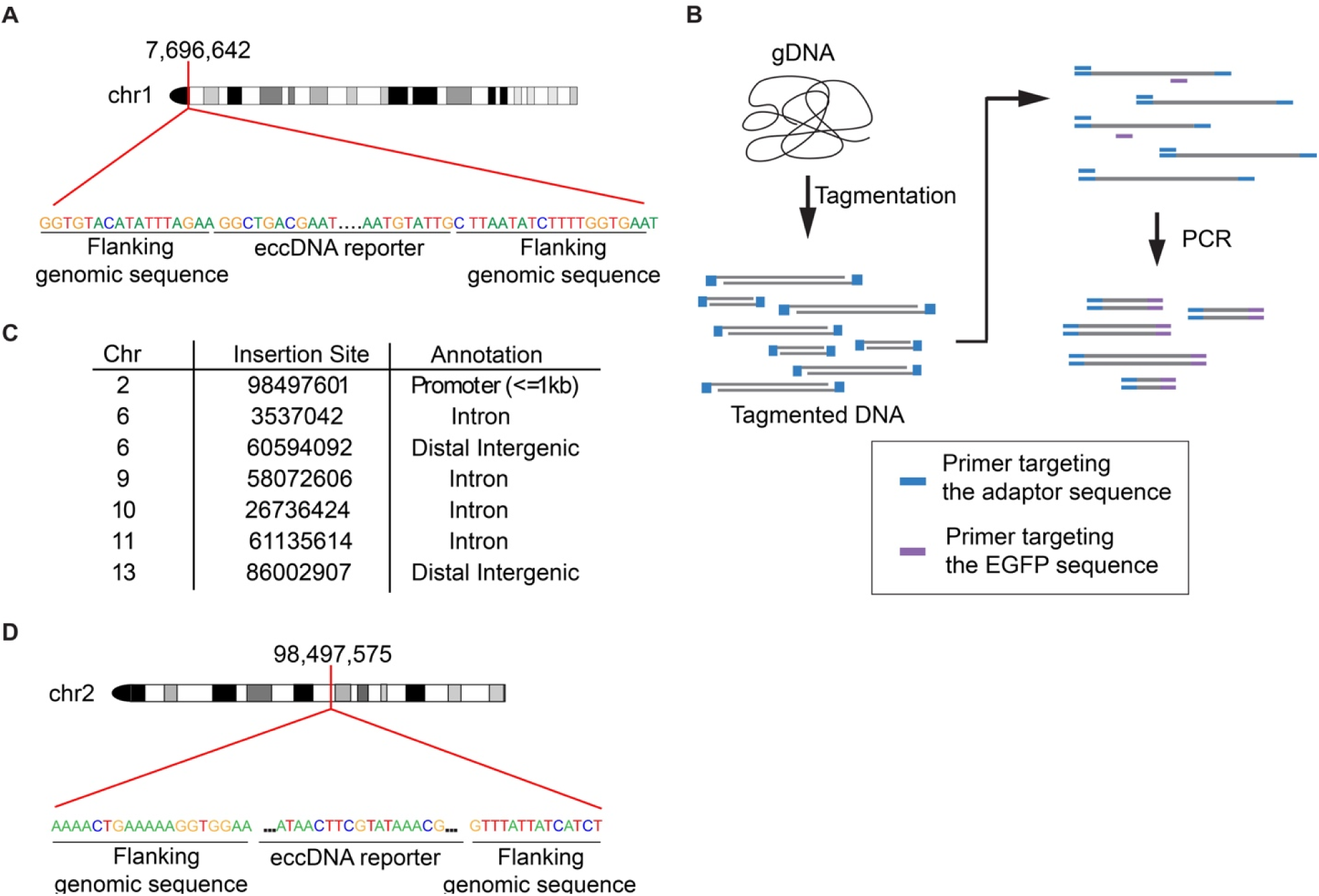
eccDNA^EGFP^ can re-integrate into the linear germline genome. (A) Whole genome long read sequencing of the genomic DNA of sperm from an AAV-Cre-injected eccDNA mouse revealed an instance of eccDNA^EGFP^ insertion to chromosome 1. (B) Schematic diagram of the targeted approach to enrich for genomic regions with eccDNA^EGFP^ insertions. (C) Annotations of eccDNA^EGFP^ genome insertion sites using the targeted long-read sequencing results of the sperm from an eccDNA mouse injected with AAV-Cre. (D) Targeted long read sequencing of the genomic DNA of sperm from CRISPR-C-treated eccDNA mouse revealed an instance of eccDNA^EGFP^ insertion to chromosome 2.

**Figure S3.**
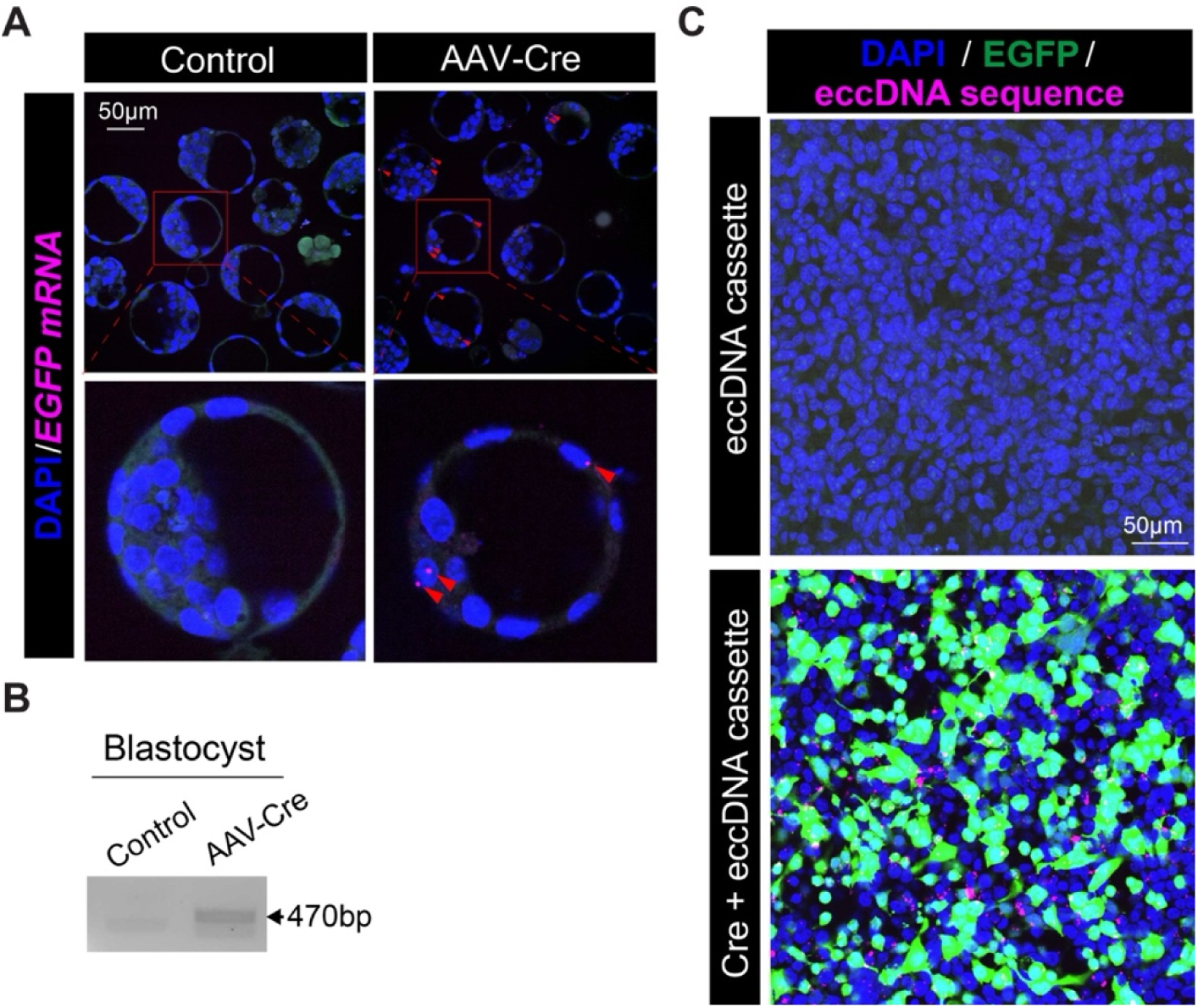
Male germline eccDNA sequences are inheritable. (A) Representative RNA FISH images of blastocyst embryos. Red arrow heads denote EGFP mRNA. (B) Positive PCR signals of eccDNA^EGFP^ junction sequence were identified in the DNA of blastocyst embryos generated using sperm from the AAV-Cre injected eccDNA mice. (C) Representative images of the RCA-based DNA FISH experiment on the HEK293T cells transfected with the reporter eccDNA components.

**Figure S4.**
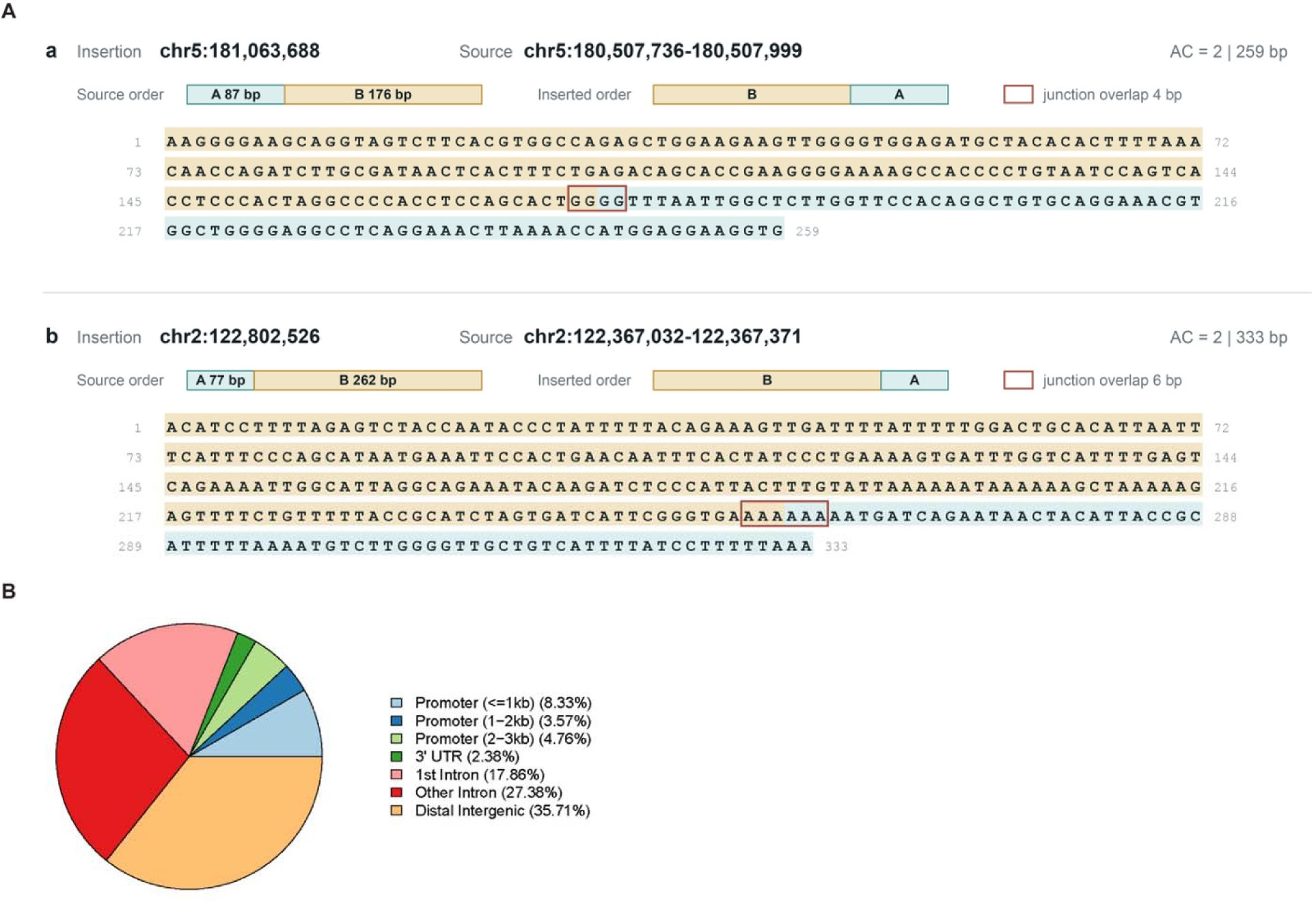
eccDNA can integrate into the human germline genome. (A) Two representative examples of eccDNA-mediated insertions in the human genome. (B) Annotations of the eccDNA genome insertion sites.

**Figure S5.**
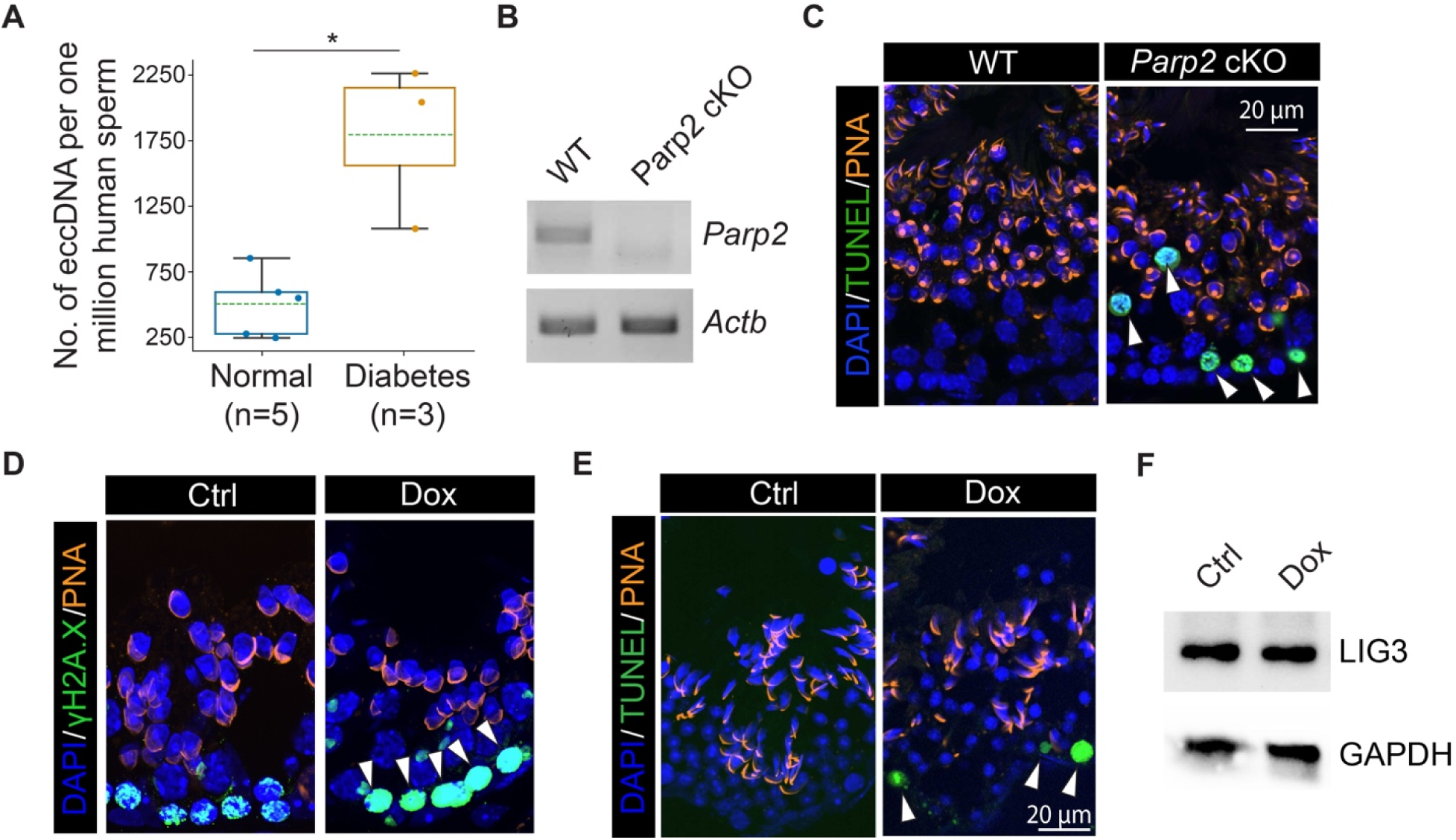
The cell signaling pathways were significantly altered in the germline of diabetic males. (A) Bar graph showing the number of sperm eccDNA in individuals with different health status. *p* values were calculated using a two-tailed Mann-Whitney U tests. *, *p* < 0.05. (B) PCR on the *Parp2* cKO germ cell cDNA confirmed the deletion of *Parp2* exon 8. (C) Representative images of the seminiferous epithelium of the WT and *Parp2* cKO testis. The TUNEL assay signals are in green (white arrow heads). The acrosomes were visualized with peanut agglutinin (PNA). (D) Representative images of the seminiferous epithelium of the control and doxorubicin (Dox)-injected testis. The γH2A.X signals are in green. White arrow heads denote increased DBS signals in the germ cells of the Dox-treated testis. The acrosomes were visualized with PNA. (E) Representative images of the seminiferous epithelium of the control and Dox-injected testis. The TUNEL assay signals are in green (white arrow heads). The acrosomes were visualized with PNA.

**Figure S6.**
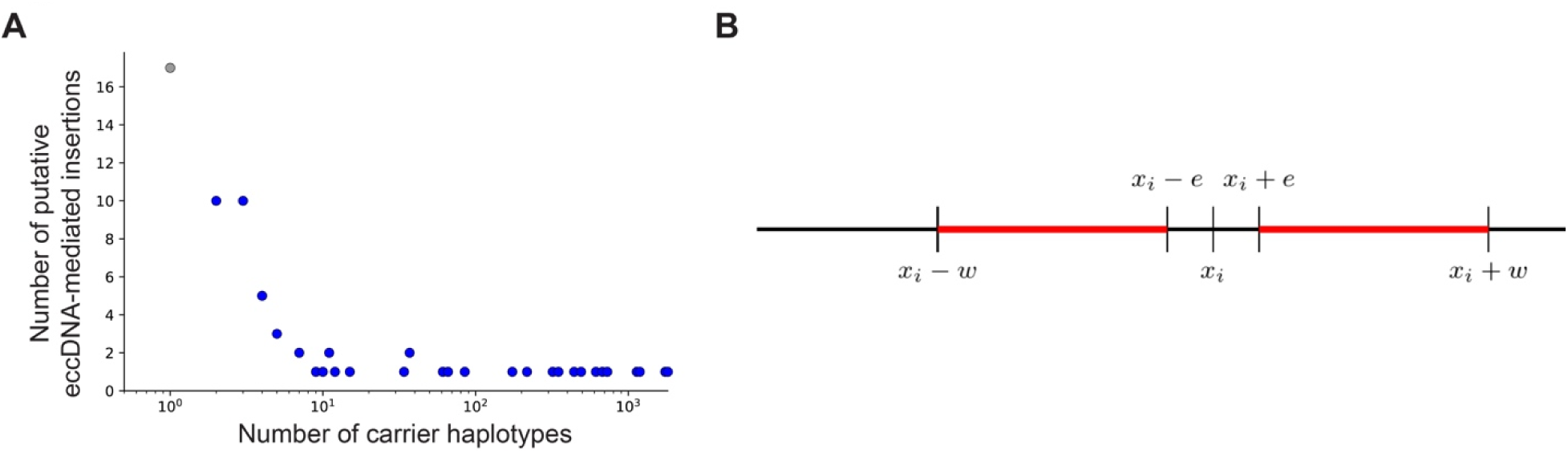
The method to estimate the local relatedness around shared eccDNA-mediated insertions. (A) Distribution of the putative eccDNA-mediated genome insertion allele counts. The gray point marks insertions observed in single carriers, while blue dots show insertions observed in two or more carrier haplotypes which passed quality controls. (B) Graphical representation of the flanking interval over which coalescent trees are extracted from the ARG and used to obtain.

## Notes

### Competing Interest Statement

The authors have declared no competing interest.

